# Breakthrough infections by SARS-CoV-2 variants boost cross-reactive hybrid immune responses in mRNA-vaccinated Golden Syrian Hamsters

**DOI:** 10.1101/2023.05.22.541294

**Authors:** Juan García-Bernalt Diego, Gagandeep Singh, Sonia Jangra, Kim Handrejk, Manon Laporte, Lauren A. Chang, Sara S. El Zahed, Lars Pache, Max W. Chang, Prajakta Warang, Sadaf Aslam, Ignacio Mena, Brett T. Webb, Christopher Benner, Adolfo García-Sastre, Michael Schotsaert

**Affiliations:** Infectious and Tropical Diseases Research Group (e-INTRO), Biomedical Research Institute of Salamanca-Research Centre for Tropical Diseases at the University of Salamanca (IBSAL-CIETUS), Faculty of Pharmacy, University of Salamanca, Spain; Department of Microbiology, Icahn School of Medicine at Mount Sinai New York, NY, USA; Global Health and Emerging Pathogens Institute, Icahn School of Medicine at Mount Sinai New York, NY, USA; Graduate School of Biomedical Sciences, Icahn School of Medicine at Mount Sinai, New York, NY, USA; NCI Designated Cancer Center, Sanford-Burnham Prebys Medical Discovery Institute, 10901 N Torrey Pines Rd, La Jolla, CA 92037, USA; Department of Medicine, University of California San Diego, La Jolla, CA, USA; Department of Veterinary Sciences, University of Wyoming, Laramie, WY, USA; Department of Medicine, Division of Infectious Diseases, Icahn School of Medicine at Mount Sinai New York, NY, USA; The Tisch Cancer Institute, Icahn School of Medicine at Mount Sinai New York, NY, USA

**Keywords:** SARS-CoV-2, variants of concern, hybrid immunity, mRNA vaccines, breakthrough infection

## Abstract

Hybrid immunity to SARS-CoV-2 provides superior protection to re-infection. We performed immune profiling studies during breakthrough infections in mRNA-vaccinated hamsters to evaluate hybrid immunity induction. mRNA vaccine, BNT162b2, was dosed to induce binding antibody titers against ancestral spike, but inefficient serum virus neutralization of ancestral SARS-CoV-2 or variants of concern (VoCs). Vaccination reduced morbidity and controlled lung virus titers for ancestral virus and Alpha but allowed breakthrough infections in Beta, Delta and Mu-challenged hamsters. Vaccination primed T cell responses that were boosted by infection. Infection back-boosted neutralizing antibody responses against ancestral virus and VoCs. Hybrid immunity resulted in more cross-reactive sera. Transcriptomics post-infection reflects both vaccination status and disease course and suggests a role for interstitial macrophages in vaccine-mediated protection. Therefore, protection by vaccination, even in the absence of high titers of neutralizing antibodies in the serum, correlates with recall of broadly reactive B and T-cell responses.

## INTRODUCTION

In 2019, severe acute respiratory syndrome coronavirus 2 (SARS-CoV-2) started circulating in the human population and caused the COVID-19 pandemic, infecting millions of people worldwide. This resulted in severe economic and human losses, lockdowns, and a race against the virus to find and develop antivirals and develop new vaccines. In the past two years, several SARS-CoV-2 vaccines, such as mRNA vaccines Pfizer BNT162b2 and Moderna mRNA-1273, have been approved and administered worldwide to confer protection against severe SARS-CoV-2 disease. mRNA vaccines have been shown to produce markedly higher antibody responses than the licensed adenoviral vector vaccines, even in suboptimal dosage regimes^1^. As a result of natural infection with or without vaccination, SARS-CoV-2-specific immunity is present in large parts of the human population. Short-term effectiveness of these vaccines has been validated in multiple clinical and preclinical studies^2–5^. However, the immune responses induced by these vaccines typically target the Spike (S) surface glycoprotein of the ancestral SARS-CoV-2 strain and might be less efficient in providing protection against variants of concern (VoCs) due to variation in S protein amino acid sequence and thereby viral escape from pre-existing vaccine or infection-derived neutralizing antibodies. Although many previous reports suggest that short-term and broadly protective immune responses against different VoCs can be induced by vaccination^6^, long-term protection conferred by these SARS-CoV-2 vaccines has may not be sufficient to protect, both in terms of disease severity and viral titers, against the ever-emerging circulating SARS-CoV-2 variants due to waning antibodies and viral escape from neutralizing antibodies as we have observed for the Omicron VoCs^7^.

Breakthrough infections, a term used to describe detectable viral infections in preimmune hosts such as vaccinated individuals, have been reported since the advent of VoCs like Alpha, Beta, Delta and Omicron variants^8–11^. Even though vaccinated individuals are more protected against SARS-CoV-2 infection and show less severe disease progression or illness than unvaccinated individuals, they can still have high viral loads and transmissibility upon infection^12–14^. Moreover, immune responses in vaccinated as well as convalescent individuals wane over time, which is another reason why reinfections can occur and further illustrates the need for booster vaccine doses to increase in anti-spike IgG responses in order to tackle emerging SARS-CoV-2 variants^15, 16^. Protective host immune responses induced by natural infection or vaccination consist of humoral responses, such as neutralizing antibodies (nAbs), as well as cellular responses including CD4^+^/CD8^+^ T cell responses specific to SARS-CoV-2 antigens^17–20^. Although mRNA vaccines induce very high binding and neutralizing antibody titers and some CD8^+^ T cells and CD4^+^ T cells, infection generates more CD8^+^ T cells, critical for clearance of infected cells. A combination of humoral and cellular immune responses may lower the probability of subsequent infections, limit viral replication as well as disease progression in infected individuals^21, 22^. The term ‘hybrid immunity’ is used to describe the host immune status resulting from a combination of vaccination and natural infection, such as in the case of breakthrough infection^23^. It is well established that vaccinated individuals show higher nAbs titers and antigen-specific T-cell responses than unvaccinated individuals after SARS-CoV-2 infection^20, 24–26^. However, there are multiple reports suggesting these immune responses wane over time, supporting the importance of vaccination amidst rising breakthrough infection cases.

Antibodies induced in response to vaccination or infection are maintained in circulation by long-lived plasma cells and can target incoming virus immediately if they reach to the appropriate tissues. Additionally, these immune responses can be boosted soon after reinfection via recall of memory B and T cells, resulting in hybrid immunity, which might be important to control the virus replication and therefore, disease severity^27^. Preliminary findings in animal models, such as non-human primates, highlighted recall of antigen specific IFN-γ^+^ T cells upon re-exposure to SARS-CoV-2 S/N peptides/peptide pools, which correlated with protection from severe disease^28, 29^. This is especially important as nAbs titers decline over time after vaccination or previous exposure and thus SARS-CoV-2-specific CD4^+^ and CD8^+^ T cells may confer rapid protection against the virus in the absence of a potent humoral response^30^. Recall kinetics highly depend on the quantity and quality of the pre-existing antigen-specific memory T cell pool, which may affect duration and dynamics of immune responses when re-exposed to VoCs.

Syrian golden hamsters are highly susceptible to SARS-CoV-2 infection showing a disease phenotype that resembles disease observed in human COVID-19 cases. Thus, to study the induction of hybrid immunity in vaccinated hosts during breakthrough infection by ancestral SARS-CoV-2 and VoCs, we performed immune profiling assays as well as an in-depth analysis of the host transcriptome using the Syrian golden hamster model for COVID-19 in naive and vaccinated animals.

## RESULTS

### Suboptimal vaccination leads to binding antibody titers against ancestral spike protein but not virus neutralization antibody titers

Following a suboptimal vaccination strategy, consisting of a single 5 µg dose of Pfizer BNT162b2 vaccine, the SARS-CoV-2 Spike (S)-specific IgG binding and neutralization capacity against the different variants was evaluated from sera collected three weeks post immunization in both vaccinated and unvaccinated animals (Figure 1). All 30 hamsters vaccinated with Pfizer-BNT162b2 showed ancestral trimeric full-length S-specific total IgG ELISA titers, confirming successful seroconversion after vaccination (Fig. 1B).

**Figure 1.**
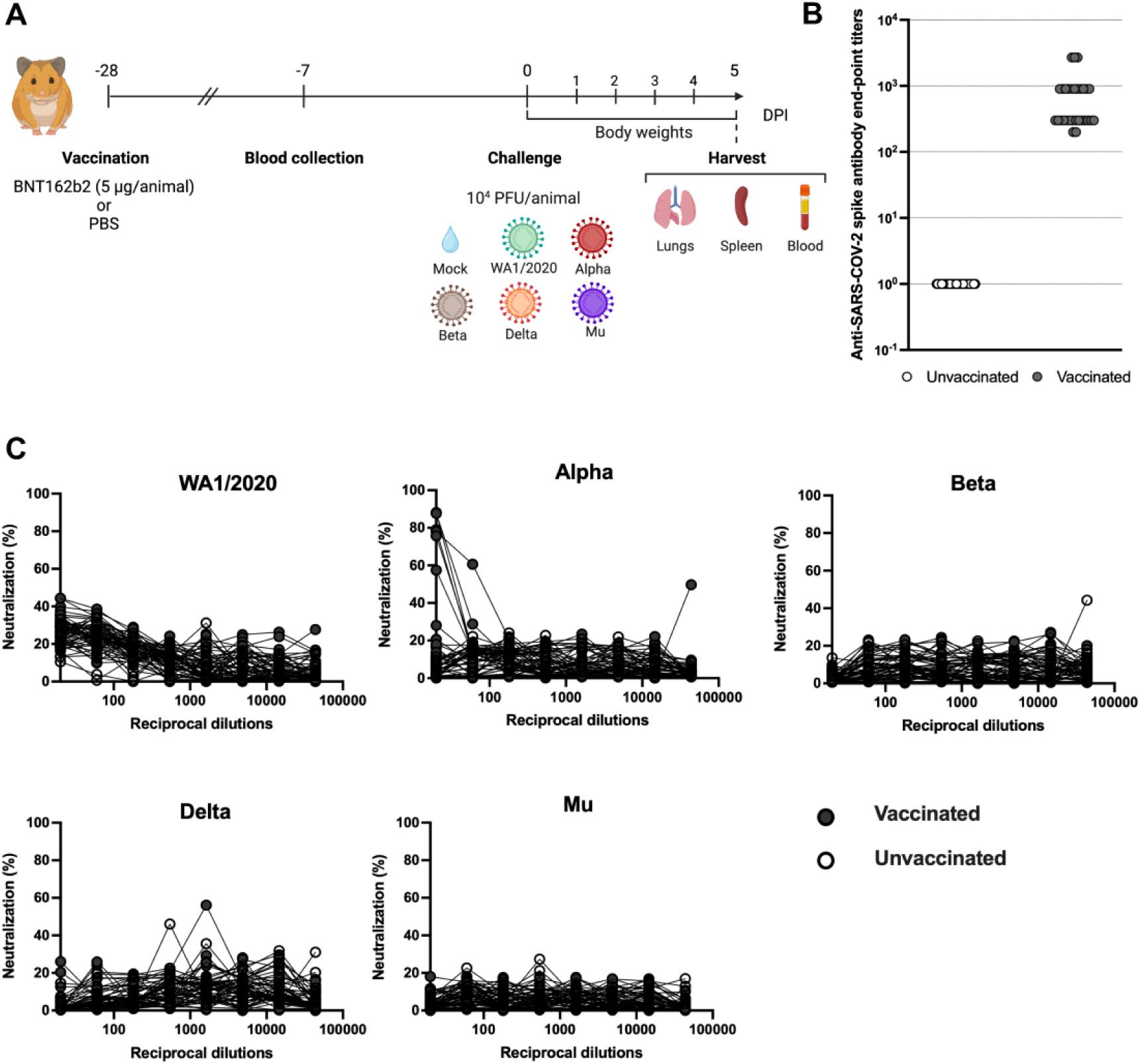
Study design and total IgG ELISA titers post vaccination. **(A)** Vaccination and SARS-CoV-2 challenge study design. Syrian golden hamsters were given 5 μg of Pfizer BNT162b2 mRNA vaccine or PBS via the intramuscular route once (n=30 hamsters/group). Blood was collected for serology 3 weeks post vaccination. Animals were either mock-challenged or challenged with the indicated SARS-CoV-2 variants. Morbidity was monitored daily as body weight changes and lungs, spleen and blood were collected at 5 days post infection (DPI). Figure created with Biorender.com **(B)** USA-WA1/2020 SARS-CoV-2 S-specific IgG ELISA titers in hamster sera 3-weeks post-vaccination. Pfizer-BNT162b2 vaccinated hamsters (n=30) are presented in grey and unvaccinated controls (n=30) are presented in white. **(C)** Microneutralization assays in hamster sera 3-weeks post vaccination against USA-WA1/2020, Alpha, Beta, Delta and Mu variants. The assay was performed with 350TCID50 per well on VeroE6/TMPRSS2 in all cases.

Next, the levels of virus-neutralizing antibodies (nAbs) were evaluated via micro neutralization assays against USA-WA1/2020 (WA1/2020 for short) and SARS-CoV-2 variants of concern (VoCs) (Fig. 1C). For ancestral WA1/2020 and Alpha, some virus neutralization activity was observed at the highest concentration of serum antibody, however this neutralization activity was absent in further dilutions. No neutralizing antibodies were detected for other VoCs, illustrating that this vaccination dose did not efficiently induce virus-neutralizing antibodies.

### Suboptimal vaccination results in faster recovery after ancestral USA-WA1/2020 and Alpha intranasal challenge but not for other VoCs

To evaluate protection against different SARS-CoV-2 VoCs, vaccinated and unvaccinated hamsters were challenged with 1×10^4^ PFU/animal of WA1/2020, Alpha, Beta, Delta, or Mu variants at 4 weeks post-immunization. Each variant was used to challenge 5 unvaccinated and 5 vaccinated animals. Two mock groups (one vaccinated and one unvaccinated) were also included in the study. From 0 to 5 days post-infection (DPI), body weights were recorded to assess the morbidity and disease severity in each group (Fig 2A-F).

**Figure 2.**
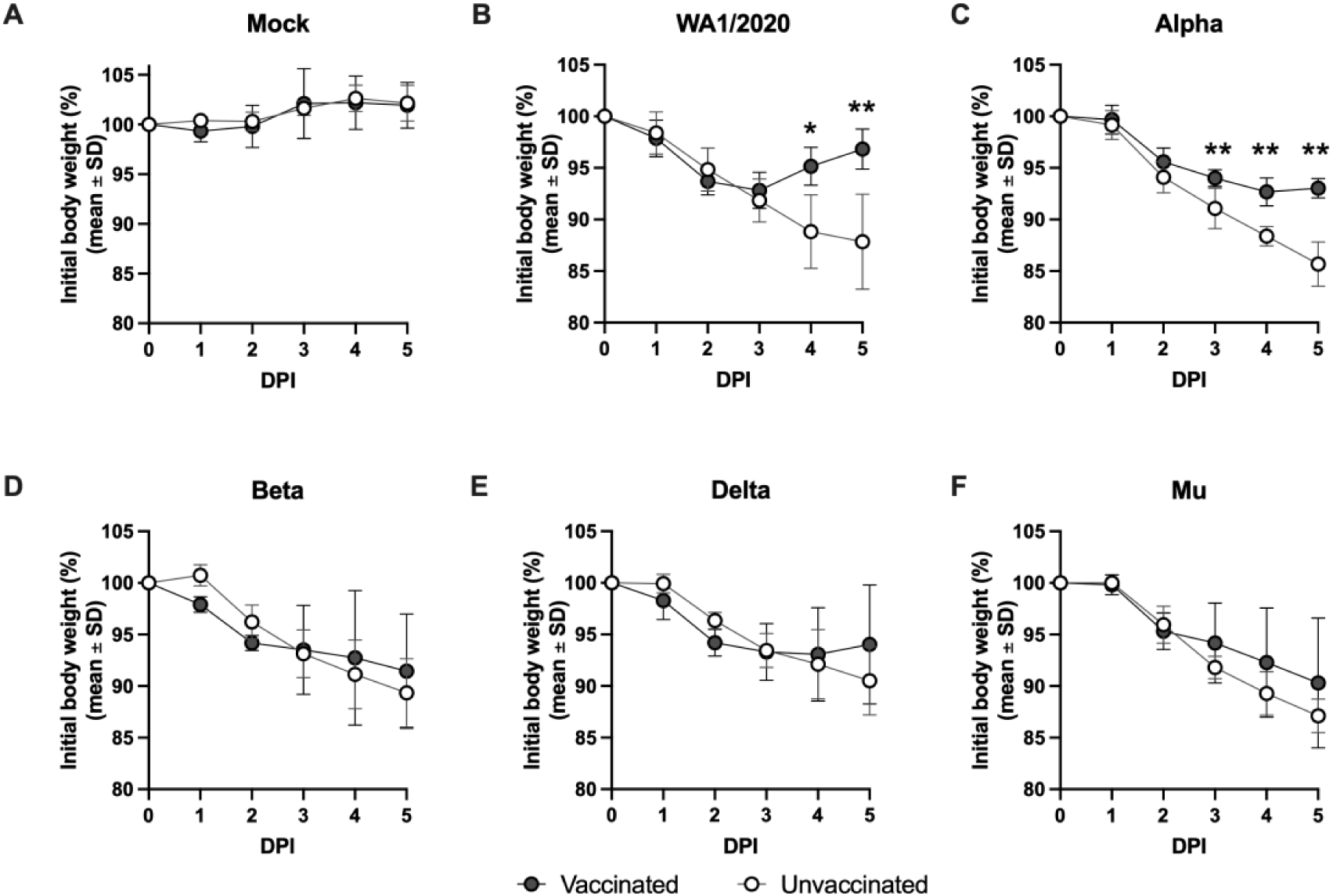
Body weight loss in hamsters with and without suboptimal BNT162b2 vaccination after challenge with USA-WA1/2020, Alpha, Beta, Delta or Mu variants. Hamster body weight measured after a challenge from 0 to 5 DPI. **(A)** Body weight loss in vaccinated (n=5) and unvaccinated (n=5) hamsters upon mock-challenge, 10^4^ PFU/animal of (**B**) ancestral WA1/2020 virus, **(C)** Alpha variant, **(D)** Beta variant, **(E)** Delta variant, or **(F)** Mu variant. Statistical analysis: n = 5/group; *p < 0.05, **p < 0.01 by Mann-Whitney U test.

SARS-CoV-2 infection in all animals resulted in body weight loss ranging from 9.5% to 14.3% by 5 DPI in the unvaccinated groups (Fig 2B-F). All SARS-CoV-2 challenged animals showed progressive body weight loss and signs of morbidity irrespective of their vaccination status starting 1 DPI (Fig. 2B-F). The BNT162b2-vaccinated groups showed faster recovery from infection compared to the respective unvaccinated groups upon challenge with WA1/2020 and Alpha variant, with the former group completely recovering to 100% body weight by 5 DPI (Fig. 2B, C). Differences in body weight loss were statistically significant between vaccinated and unvaccinated animals at 4 DPI upon challenge with either WA1/2020 or Alpha variant (Fig. 2B, C). WA1/2020-challenged vaccinated hamsters had 96.8% ± 1.9% of their initial bodyweights, while unvaccinated ones presented 87.8% ± 4.6% (Fig. 2B) at 5 DPI. Similarly, Alpha-challenged-vaccinated animals showed 93.0% ± 0.9% of their initial bodyweights, while unvaccinated animals presented 85.7% ± 2.2% (Fig. 2C).

However, in the case of Beta, Delta and Mu challenged animals, no significant differences in weight loss were found between vaccinated and unvaccinated groups. Regardless of vaccination, all animals lost body weight progressively from 1 to 5 DPI (Fig. 2D-F). For vaccinated hamsters infected with Delta, signs of slight recovery were shown at 5 DPI, presenting 94.0% ± 5.8% of their initial bodyweights while the differences with the unvaccinated group (90.5% ± 3.3%) were not statistically significant (Fig. 2E). Both Beta and Mu showed no signs of protection by vaccination with no significant differences in bodyweight at 5 DPI between vaccinated and unvaccinated groups (vaccinated: 91.4% ± 5.6% *vs.* unvaccinated: 89.3% ± 3.3% for Beta; vaccinated: 90.3% ± 6.3% *vs.* unvaccinated 87.1% ± 1.6% for Mu) (Fig. 2D and F respectively).

### Suboptimal vaccination abrogates WA1/2020, Alpha, and Delta viral titers but results in breakthrough infection with detectable virus at 5 DPI after Beta and Mu challenge

To further evaluate protection against SARS-CoV-2 and variants conferred by single dose of 5 μg Pfizer BNT162b2 vaccination, viral titers in the lung at 5 DPI were examined by plaque assay. Intranasal infection with any of the challenge viruses resulted in detectable lung viral titers at 5 DPI in all unvaccinated groups. The geometric mean titers (GMT), as measured in Vero E6/TMPRSS2 cells, ranged from 6.87×10^3^ PFU/mL for Delta to 4.68×10^5^ PFU/mL for Beta, with 1.25×10^5^ PFU/mL for WA1/2020, 1.81×10^5^ PFU/mL for Alpha and 9.79×10^4^ PFU/mL for Mu. In the case of vaccinated groups, no detectable viral load was found in the lungs of hamsters infected with WA1/2020 or Alpha variant at 5 DPI. Additionally, no detectable titers were observed in the lungs of 4 out of 5 vaccinated animals challenged with Delta. Conversely, viral titers were detectable in the lung at 5 DPI for all unvaccinated and vaccinated animals challenged with Beta (1.15×10^3^ PFU/mL) or Mu (6.22×10^2^ PFU/mL) variants. However, a significant reduction in lung viral titers were observed even with the variants implicated in breakthrough infection (Figure 3).

**Figure 3.**
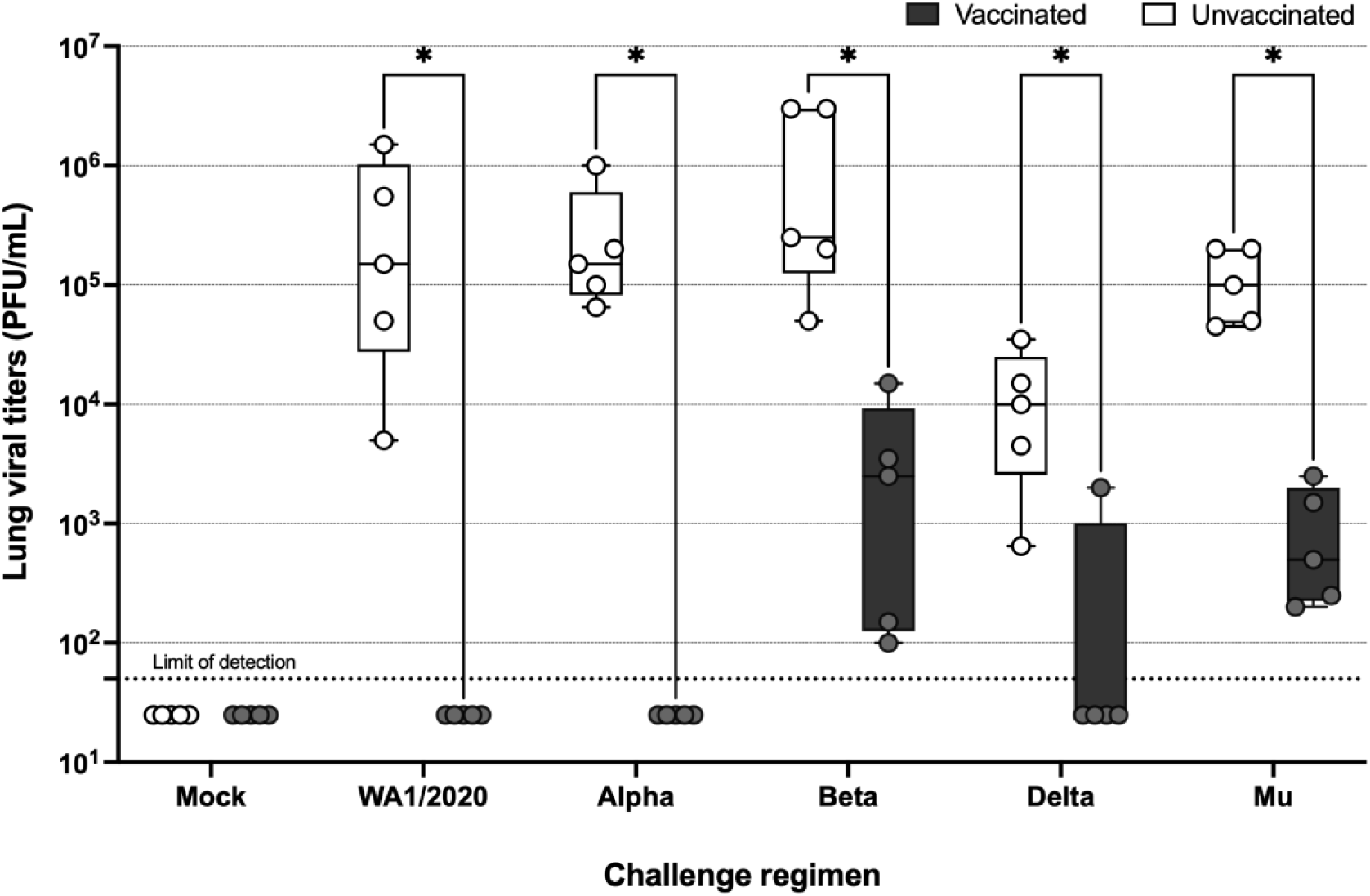
Viral titers in the lung measured via plaque assays in VeroE6/TMPRSS2 cells. Limit of detection (LOD) = 50 PFU/mL. Statistical analysis: n = 5/group; *p < 0.05 by Mann-Whitney U test.

### T cells are primed by vaccination and boosted by infection

To assess T cell activation induced by the BNT162b2 vaccination and subsequent challenge, spleens were harvested at 5 DPI to assess IFN γ^+^ cell induction using ELISpots. Activation of T cells in naïve hamsters is previously reported to be initiated by but not peak around 5 DPI^31^. Therefore, this timepoint allowed us to focus on specific T cell responses conferred by vaccination, and not by *de novo* T cell responses to infection. Splenocytes were restimulated with overlapping 15-mer peptides of either SARS-CoV-2 Spike (S), Nucleoprotein (N) or an irrelevant Hemagglutinin (HA) peptide as non-specific control. The final number of IFN γ-releasing splenocytes per million was calculated for each spleen stimulated with peptides (S, N, HA) or no stimulation (Fig 4A and Suppl Fig 1). Fold-induction as a ratio of N or S-restimulated splenocytes with irrelevant HA as a reference was also calculated (Fig 4B).

**Figure 4.**
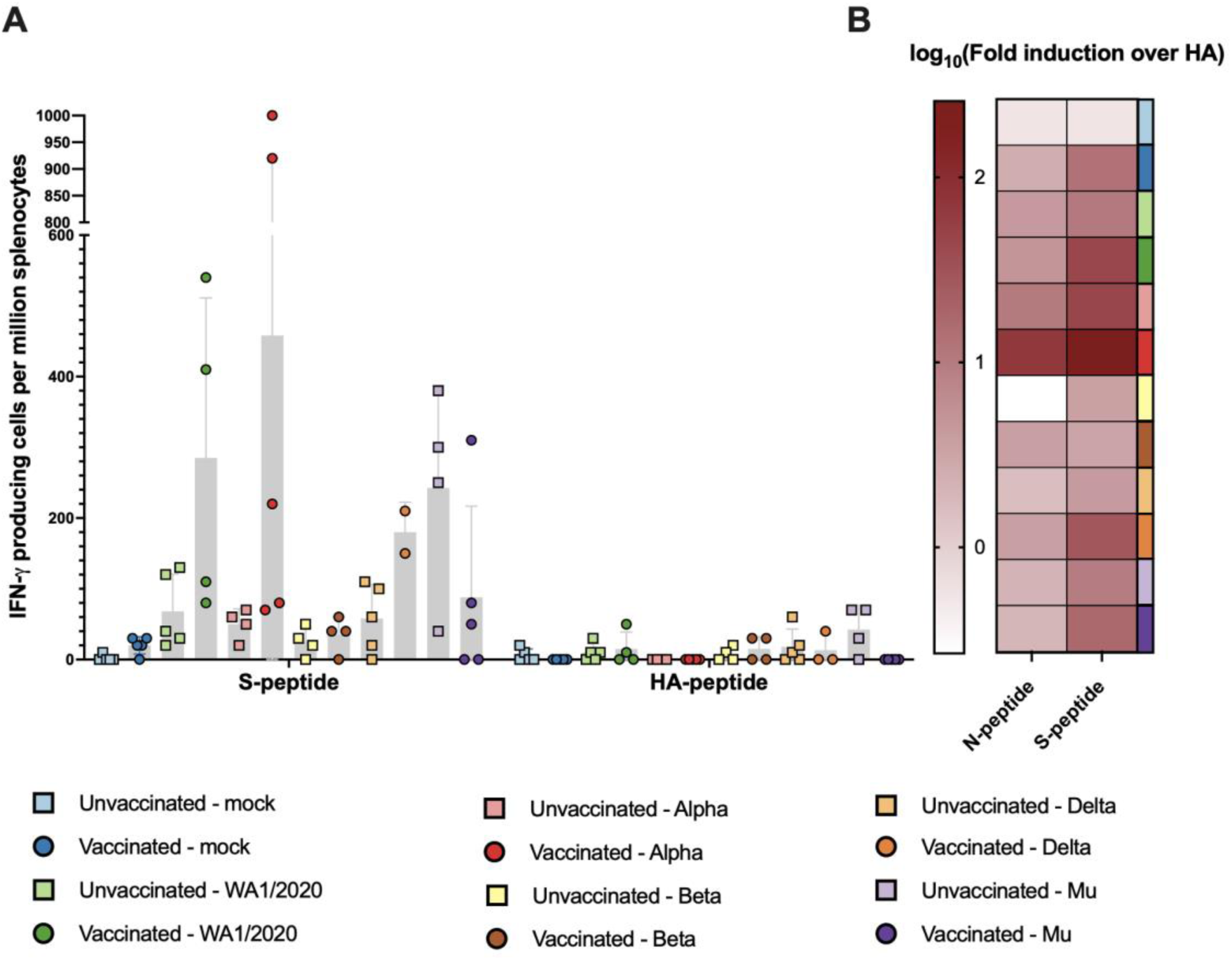
T cell activation measured by IFN-γ^+^ ELISpots from splenocytes. N-peptide: 15-mer overlapping peptides based on the Nucleoprotein (N) sequence of SARS-CoV-2. S-peptide: 15-mer overlapping peptides based on the Spike (S) sequence of SARS-CoV-2. (A) IFN-γ-releasing splenocytes per million after stimulation with S or irrelevant Hemaglutinin (HA). (B) Log_10_ of the fold induction calculated based on number of IFN-γ-releasing splenocytes when stimulated with S or N taking HA as reference.

S-specific T cell activation, measured as IFN γ-releasing cells per million splenocytes, was higher in splenocytes harvested from all the vaccinated animals as compared to unvaccinated animals in respective groups, except groups challenged with Beta or Mu (Fig 4A). Additionally, limited NP-specific T cell activation was detected, although it was also higher in vaccinated groups compared to the respective unvaccinated group (Supplementary Figure 1). In all, these results suggest that BNT162b2 vaccination primed T cell responses that can be boosted by subsequent virus challenge. Fold activation was higher in splenocytes from vaccinated animals infected with either WA1/2020 virus (mean fold induction = 185) and Alpha variant (mean fold induction = 458), correlating with more protection from body weight loss and the highest reduction lung viral titers. Similarly, activation of S specific T cells was also observed in animals challenged with Delta (mean fold induction = 106.88) and Mu variants (mean fold induction = 88.65) but to a lesser extent as compared to WA1/2020 and Alpha variant. Mu-challenged animals also had high spike-specific T cell induction in unvaccinated animals, but also showed higher activated T cell levels upon HA-peptide restimulation. Therefore, when fold induction over HA is calculated for the groups challenged with the Mu variant, higher fold induction is seen for the vaccinated group, as compared to unvaccinated animals. In the case of the Beta variant, although some fold induction can be measured (mean = 15.92), almost negligible T cell activation was detected (Fig. 4B)

### Infections with VOCs in suboptimal vaccination broadens antibody protection against both VOC and ancestral WA1/2020

Next, we evaluated the breadth of serum neutralization capacity against ancestral virus and all of the different challenge VoCs using microneutralization assays. Neutralization of the WA1/2020 strain was observed in all the vaccinated groups, regardless the VoC used for challenge. Additionally, neutralization of the WA1/2020 virus was observed even in unvaccinated animals after challenge with either WA1/2020, Alpha variant and, to a lesser extent, Delta variant. This suggested that infection is more effective at inducing nAb activity than a single dose of BNT162b2. Nevertheless, no WA1/2020 neutralization capacity was detected in the serum of animals challenged with antigenically more distant Beta or Mu variants (Fig. 5A and B).

**Figure 5.**
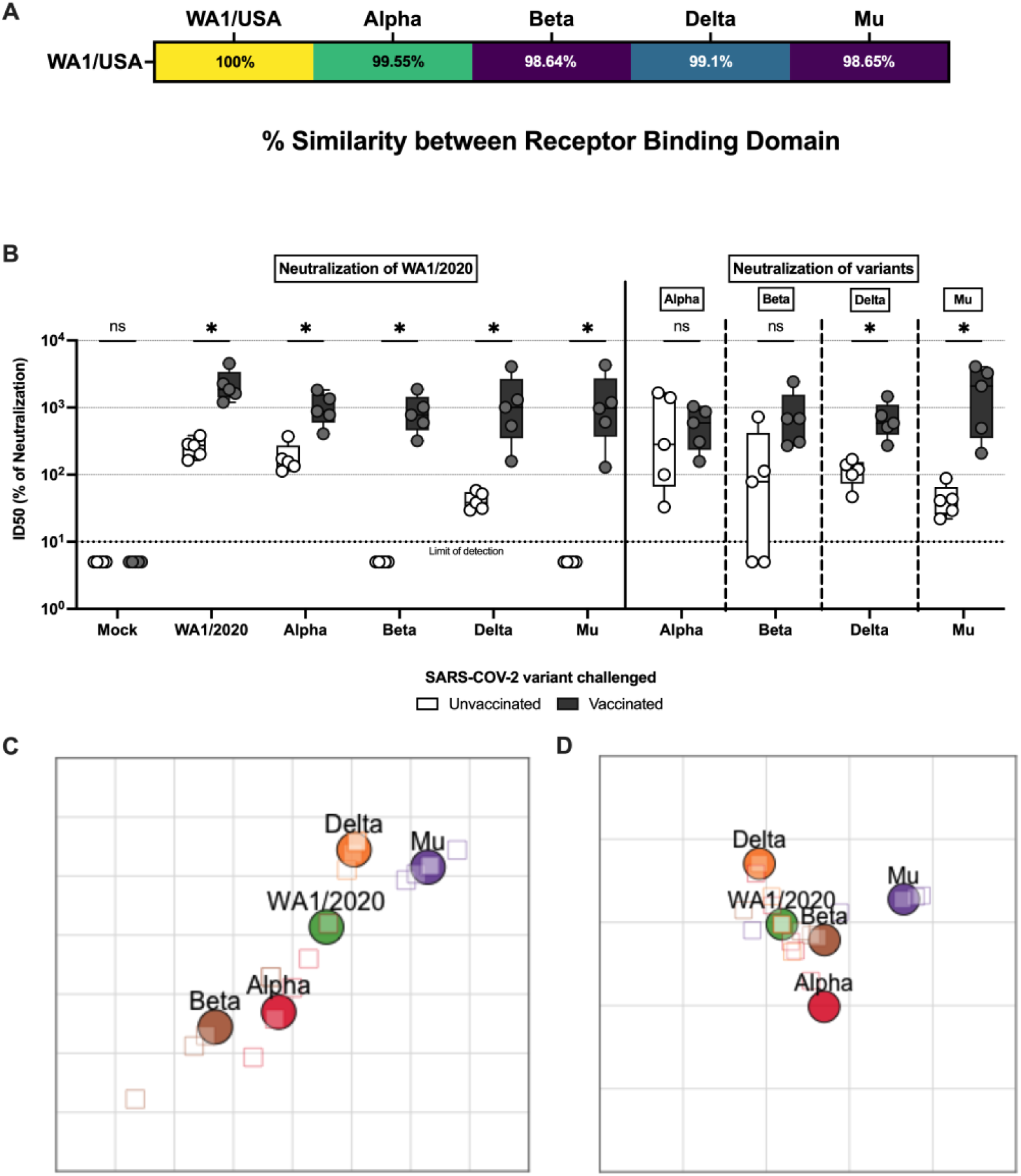
Antibody neutralization activity after challenge. **(A)** Similarity of the receptor binding domain (RBD) based on amino acid sequence of Alpha, Beta, Delta, and Mu compared with ancestral WA1/2020 expressed as the similarity percentage. **(B)** Micro-neutralization assays performed in the presence of 350 TCID_50_ per well of virus and sera collected 5 DPI. In all cases, the left panel represents neutralization activity against USA-WA1/2020 (clear: unvaccinated; solid: vaccinated) and the right panel neutralization activity against the variant of infection (clear: unvaccinated; solid: vaccinated). Statistical analysis: n = 5/group; *p < 0.05 by Mann-Whitney U test. **(C)** Antigenic map constructed with ID50 values of unvaccinated animals. Each square in the grid represents one antigenic distance unit **(D)** Antigenic map constructed with ID50 values of BNT162b2 primed animals. Each square in the grid represents one antigenic distance unit.

In the case of Alpha, neutralizing capacity against both WA1/2020 and Alpha were comparable in vaccinated animals, and potent neutralizing activity against both was also observed in unvaccinated infected animals. A similar pattern can be observed in hamsters infected with Delta, although the neutralization capacity in the case of unvaccinated animals is reduced, especially against WA1/2020 (Fig. 5B).

Animals infected with Beta or Mu show a similar antibody response. In both cases, vaccinated animals present high antibody neutralization activity against both WA1/2020 and Beta or Mu, respectively. Unvaccinated animals present some neutralization capacity against the variant used for infection, although reduced in comparison with other variants. No neutralizing antibody induction is observed against the WA1/2020 virus (Fig. 5B).

In summary, hybrid immunity induced by BNT162b2 vaccination followed by infection results in broad antibody neutralization capacity against both WA1/2020 and the variant of infection. However, infection without vaccination, results in broad antibody neutralization only when it is caused by Alpha or, to a lesser extent Delta, but it is specific to the variant of infection when it is caused by Beta or Mu.

This is further supported by the antigenic distances as measured with antigenic cartography. Antigenic distance is broadly defined as the property of two antigens where the shorter antigenic distance between them the greater number of antibodies that will be able to bind both. Antigenic maps generated with ID50 values (indirect representation of microneutralization titers), show reduced antigenic distances in vaccinated groups when compared to unvaccinated ones, supporting prior findings. Antigenic distances between Beta, Delta and Mu variants and the rest were reduced after prime with BNT162b2 (Fig 5D) compared with animals infected without BNT162b2 vaccination (Fig 5C). Conversely, antigenic distance between closely related ancestral WA1/2020 and Alpha was maintained regardless of the vaccination status.

### Suboptimal vaccination reduces lung pathology in WA1/2020 infection but has little to no effect after infections with VoCs

To evaluate the effect of infections with different VoCs on lung lesions, the left lung lobe was harvested at 5 DPI from each animal and fixed in formaldehyde for blinded histopathology scoring. Overall pulmonary lesions were mild both in vaccinated and unvaccinated animals (Fig 6A, B). General changes were indicative of suppurative and histiocytic bronchopneumonia to broncho-interstitial pneumonia with vasculitis and variable type II pneumocyte hyperplasia. Vasculitis and perivasculitis of medium caliber veins and to a lesser extent arteries were significant and consistent amongst most animals. Nevertheless, microthrombi were not detected and thus not scored. Bronchiole epithelial injury was minimal although it might be masked by reparative hyperplasia (Fig 6B). Following infection with WA1/2020, vaccinated animals showed an aparent reduction in pathology scores compared to unvaccinated hamsters (total scaled score of 3.28 unvaccinated *vs.* 1.3 vaccinated). Perivascular inflammation, vessel injuries, perivascular inflammation and type II pneumocyte hyperplasia showed lower scores in vaccinated animals. Furthermore, alveolar necrosis and bronchial necrosis scores were comparable to those of uninfected animals. Animal vaccinated with Delta variant also showed an overall, albeit modest, reduced pathology score compared to unvaccinated (from 2.5 to 2.1 total scaled score), with reduced alveolar necrosis, perivascular inflammation and vasculitis. Vaccinated animals infected with Alpha did not show a reduction in overall pathological score after vaccination (2.2 unvaccinated vs. 2.55 vaccinated). However, that scored might be skewed by the high levels of type II pneumocyte hyperplasia, associated with healing, in the vaccinated group (3.0 in vaccinated vs. 1.2 in unvaccinated. Beta (2.2 unvaccinated vs 2.3 vaccinated) and Mu (2.4 for both groups) variants obtained similar pathology scores compared to those of unvaccinated individuals (Fig 6A).

**Figure 6.**
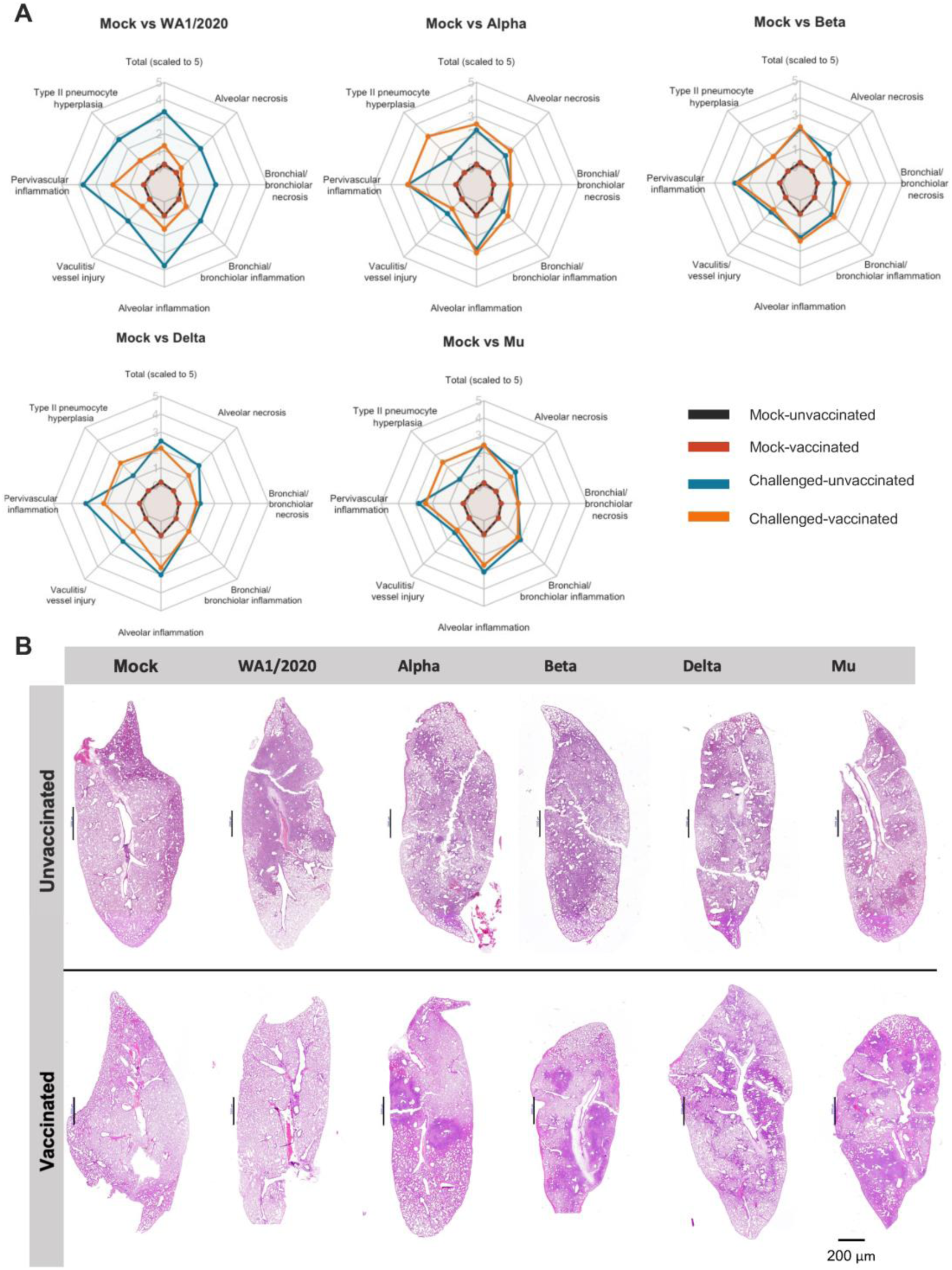
Lung pathology. **(A)** Radar charts representing mean pathology scores, scaled 1 to 5. In all cases, scores for uninfected unvaccinated (black) and vaccinated (red) individuals are included for comparison. Mean pathology scores of challenged unvaccinated individuals are shown in blue and challenged BNT162b2 primed individuals in orange. From left to right and top to bottom: Mock vs WA1/2020. Mock vs Alpha. Mock vs Beta. Mock vs Delta. Mock vs Mu. Histological parameters: 0=none, 1=minimal, 2=mild, 3=moderate, 4=marked, 5=severe. Overall lesion scores are scaled to 0-5 for representation. **(B)** Section slides with Hematoxylin and Eosin staining of lungs from one animal in each experimental group.

### Bulk RNA-seq of the lung suggests a broader immune response in groups challenged with WA1/2020 and Alpha variants than Beta, Delta and Mu

To assess how BNT162b2 prime and VoC challenge affected overall gene expression in the lung, the right lung caudal lobe was harvested from the animals at 5 DPI for bulk RNA-seq (Fig. 7). Principal component analysis (PCA) analysis shows agreement with prior findings: highest variance can be found between vaccinated and unvaccinated groups challenged with WA1/2020 and Alpha variants, suggesting a major effect of BNT162b2 vaccination on the lung response to challenge. This is particularly obvious for the groups challenged WA1/2020, for which vaccination shifts the group towards the mock. Smaller effects of vaccination are found between vaccinated and unvaccinated groups challenged with Beta, Delta, or Mu variants (Fig. 7A). Pairwise comparisons confirm an increased effect magnitude in gene expression of vaccination when followed by a challenge with WA1/2020 and Alpha variants. Alpha shows the most variation in gene expression levels when comparing vaccinated and unvaccinated groups (541 genes upregulated and 762 downregulated) followed by WA1/2020 (159 up and 371 down), Mu (131 up and 285 down) and Delta (66 up and 206 down). Almost no change in differential gene expression is detected between vaccinated and unvaccinated groups when challenged with Beta (11 up and 6 down) suggesting lower vaccination effect when challenged with this antigenically distant virus and in line with the lower antibody and T cell responses discussed in previous sections (Fig. 7B). This analysis present the limitation that although the differentially expressed genes are driven by biological differences, they can also be affected by variability between the samples. The different challenged groups showed substantial variability, that might have affected some more than others.

**Figure 7.**
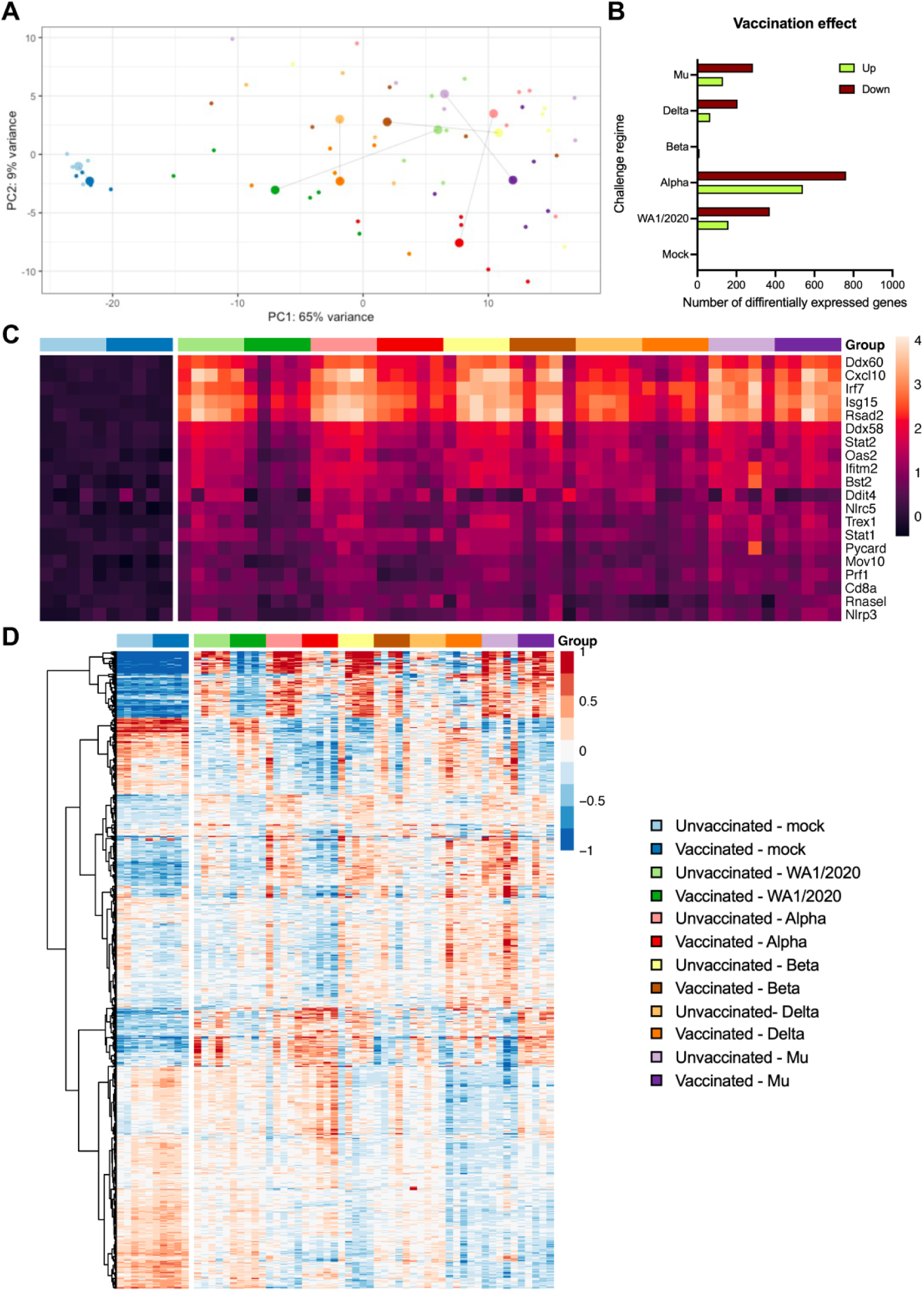
Lung RNA-seq. **(A)** PCA plot showing the relationship between vaccinated and unvaccinated groups challenged with different variants. For each group, centroids are calculated. **(B)** Pairwise comparisons of the number of up and down-regulated genes between unvaccinated and vaccinated animals challenged with each variant (adj. p<0.05). **(C)** Heatmap showing gene expression of the top 20 genes involved in the defense response to virus (GO:0051607). **(D)** Heatmap of the 1,000 most significant genes. A common legend for A, C and D panels is included.

Changes in gene expression show clear activation of viral defense mechanisms in all challenged groups when compared to mock groups (GO:0051607). Unvaccinated groups challenged with ancestral WA1/2020, Alpha, or Delta variants show higher activation of defense genes than vaccinated groups. This pattern of expression is not maintained comparing vaccinated and unvaccinated groups challenged with Beta or Mu variants (Fig 7C), showing high activation in both groups. Viral defense genes that show the highest activation include several genes involved in the regulation type I interferon (IFN) genes (IFN-α and IFN-β) and IFN-stimulated genes (ISG), such as *Ddx60*, *Irf7* or *Rsad2*; pro inflammatory cytokines such as *Cxcl10* or genes coding for ubiquitin-like proteins such as *Isg15*. Gene ontology analysis reveals a similar pattern of activation of other biological pathways that play an important role in antiviral defense including: response to other organisms (GO:0051707), biological process involved in interspecies interaction (GO:0044419), immune response (GO:0006955), immune system process (GO:0002376), response to cytokines (GO:0034097) or innate immune response (GO:0045087). Those pathways are highly upregulated in unvaccinated animals challenged with WA1/2020 or Alpha when compared to vaccinated groups. However, animals challenged with Beta, Delta or Mu, show similar gene activation pattern regardless of their vaccination status (Supplementary Figure 2).

Finally, genes upregulated in vaccinated animals infected with vaccine-matching WA1/2020 were compared to mock-infected vaccinated ones. The list was refined, removing those upregulated in unvaccinated hamsters challenged with WA1/2020 when compared to the mock-infected unvaccinated. Thus, genes showing upregulation related only to vaccination but not to infection were selected and can be found in Supplementary Table 1. Within that gene set, several show involvement in the regulation of T cells, memory T cells and NK cells, including *Zfp683* or *Klrg1*; *Zc3h12d* in antigen presentation; in the maturation and proliferation of B cells, such as *Icosl*, *Lime1*, *Cd37* or *Gpr183*; *Zc3h12d* in macrophage activation or *Nup85* in monocyte chemotaxis. Several genes related to DNA repair/replication (*Ddx11*, *Wdr76*, *Rfc3* or *Paxip1*) and genes coding for calcium-dependent proteins (*Itln1*, *Lime1*, *Themis* or *Esyt1*) were also upregulated (Supplementary Table 1).

### Immune cell abundance estimation in infected lungs using gene expression data confirm observed vaccine responses and suggest a protective role for interstitial macrophages and eosinophils

To estimate the immune cell populations in the lungs of the animals, deconvolution was performed with CIBERSORTx was performed (Fig. 8), using a signature matrix derived from a single-cell Syrian golden Hamster lung dataset^31^. All estimated cell fractions are shown in Supplementary Figure 3.

**Figure 8.**
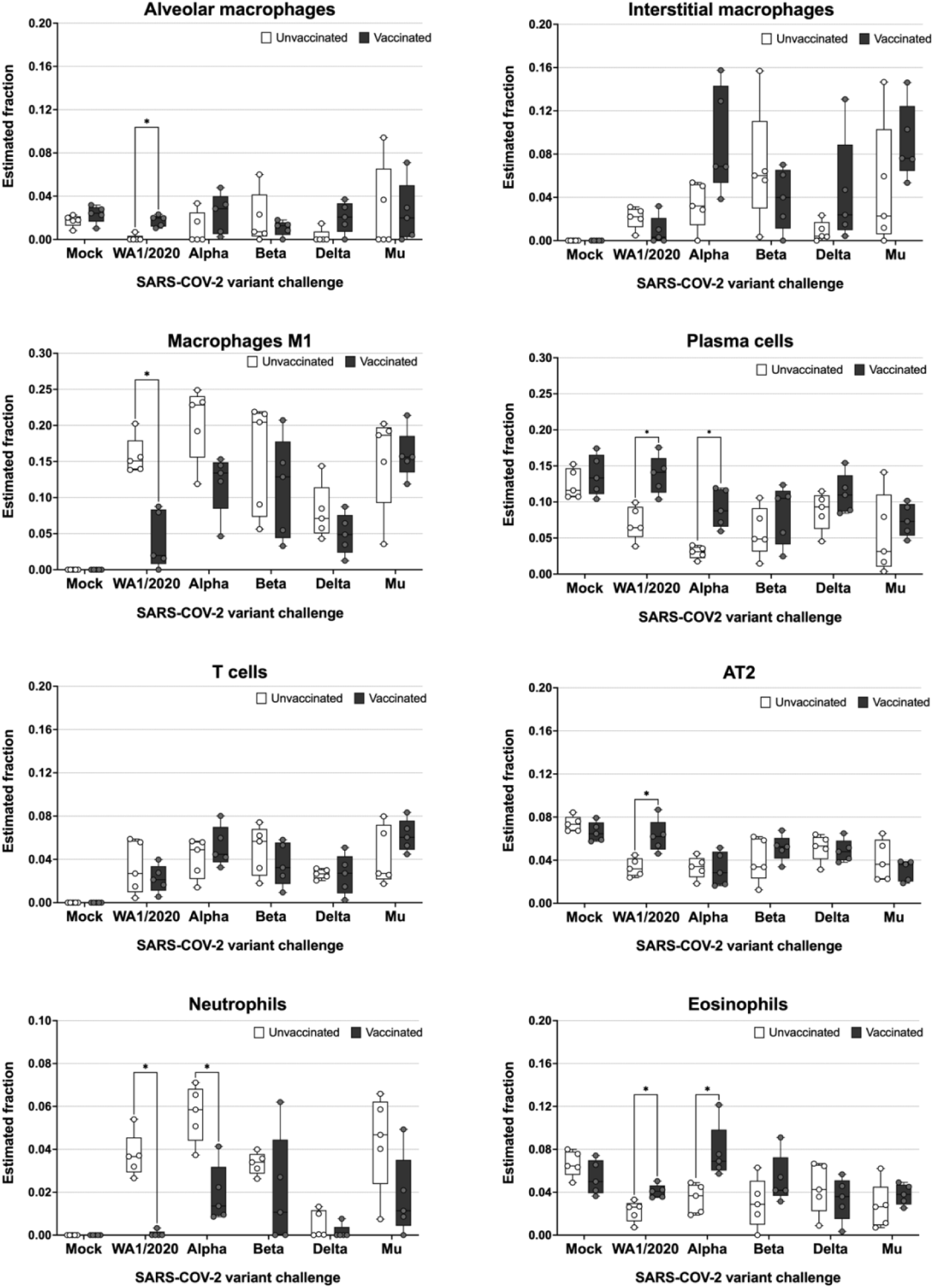
Abundance of different cell subsets in the lungs of infected hamsters extrapolated from the host transcriptome. RNA was extracted from lungs of infected hamsters at 5 DPI and subjected to bulk RNA-seq analysis. Host transcriptomes were used to impute gene expression profiles using the Cibersortx algorithm. sStatistical analysis: n = 5/group; *p < 0.05 by Mann-Whitney U test.

Alveolar macrophages seemed to be present in lower frequencies by 5 DPI for unvaccinated animals, whereas vaccination appeared to allow animals to maintain alveolar macrophage frequencies similar to that of mock-infected animals. Interstitial macrophages show elevated fractions after challenge with VoCs in vaccinated groups when compared to their respective unvaccinated groups, except for groups challenged with Beta, which show no differences. This reflected on the T cell activation profiles after challenge as described in Fig. 4. The estimated fraction of M1 macrophages in the lung was statistically significantly lower in animals vaccinated and challenged with WA1/2020 and markedly reduced when challenged with Alpha variant. When challenged with other variants, M1 levels in vaccinated animals were higher, but not statistically significantly different from unvaccinated animals.

Estimated fractions of plasma cells show a statistically significant increase in vaccinated groups challenged with WA1/2020 and Alpha variants compared to unvaccinated animals. These estimations are in line with the improved antibody response as part of hybrid immunity after infection and discussed in previous sections. Vaccination followed by infection also resulted in an increase in plasma cells upon infection with the other VoCs tested, however this increase was not statistically significant. On the other hand, lung T cells represent elevated fractions in animals challenged when compared with mock groups. Nevertheless, no differences amongst vaccinated and unvaccinated animals are detected.

Alveolar type II (AT2) cells, a cell type readily infected by SARS-CoV-2^32^, show a marked decrease at 5 DPI in all groups compared to the mock. Nevertheless, vaccination with BNT162b2 almost completely recovers AT2 levels when challenged with the ancestral WA1/2020. This suggests a protection of infection for this cell type provided by antigenically matching vaccine only for the ancestral virus.

An elevated fraction of neutrophils, associated with more severe presentations of COVID 19, are detected in unvaccinated groups challenged with all virus strains tested except for Delta. Vaccination seemed to reduce estimated neutrophil levels, suggesting vaccine mediated protection from disease. Finally, eosinophil subsets are relatively more abundant in vaccinated animals challenged with ancestral virus, Alpha, or Beta when compared to unvaccinated controls.

## DISCUSSION

Circulating SARS-CoV-2 VoCs can escape neutralizing antibodies induced by vaccination with vaccines based on ancestral spike proteins or previous infection due to the acquisition of mutations in antigenic sites in newly emerging virus variants. However, vaccination can induce immune responses that contribute to protection beyond virus neutralization, for example by the induction of non-neutralizing antibodies that can bind viral spike proteins and T cells. It is expected that host immune responses during (breakthrough) infections will be skewed and eventually boosted depending on pre existing immunity. To study the induction of hybrid immunity, the result of vaccination and natural infection, we tested host immune responses during infection with ancestral SARS-CoV-2 as well as VoCs in naive and suboptimally mRNA-vaccinated animals. This reflects the situation in the human population during the COVID-19 pandemic when mRNA vaccines became available under emergency use approval. It was observed that single vaccination already protected from severe COVID-19 even before the onset of neutralizing antibody titers in humans^5, 33^.

We show that, although limited, suboptimal vaccination in Syrian golden hamsters produces ancestral S-binding IgG titers. Nevertheless, those titers do not result in significant neutralization against the ancestral WA1/2020 virus or any of the VoCs tested. Reduced neutralization against the different VOCs is well defined in humans, with more marked decreases in neutralization of Beta, Mu, and Omicron variants, to a lesser extent of Delta, and a very slight reduction for Alpha in BNT162b2-vaccinated individuals^34–36^. Weak neutralization responses are even described after induction of high RBD and S binding antibodies after a single dose of BNT162b2 vaccine in naïve individuals^37^. The non-concordance between binding and neutralizing antibody titers suggests alternative mechanisms of immune protection from severe disease, which we observed in this study. In our study, all unvaccinated animals show a marked bodyweight reduction until 5 DPI after challenge, which marks the end of the experiment. BNT162b2 suboptimal vaccination protects against major body weight loss after challenge with the ancestral WA1/2020 virus and the antigenically similar Alpha variant. Animals either completely regain all the weight lost by 5 DPI (WA1/2020) or stop losing weight after 3 DPI (Alpha). Vaccinated hamsters challenged with Delta show a delayed recovery by 5 DPI, although not statistically significant, while vaccinated groups challenged with Beta or Mu keep losing weight by 5 DPI, illustrating that recovery after infection in vaccinated animals correlates better with antigenic match between challenge virus and vaccine. Syrian golden hamsters are highly susceptible to SARS-CoV-2 infection showing a disease phenotype that resembles the one in human COVID-19 cases. Loss of 10–20% of the initial body weight is expected 6–7 days after infection for this inoculum dose, with slight variations depending on age, variant and virus dose^38^. Maximum weight loss occurred at 3-4 DPI in the vaccinated groups that showed protection, which correlated with peak viral titers in the upper and lower respiratory tract^39^. Thus, we can conclude that suboptimal vaccination protects against WA1/2020 and Alpha, even in the absence of detectable nAb post-vaccination. Protection might be mediated by a combination of non-measurable nAb responses *ex vivo* and T cells. Body weight loss and recovery are reflected by complete control of virus in the lungs of vaccinated animals challenged with WA1/2020 and Alpha, as well as in 4 of 5 vaccinated animals challenged with Delta by 5 DPI. Although breakthrough virus is not completely cleared by 5 DPI in vaccinated groups challenged with Beta and Mu, lung titers are substantially reduced compared to unvaccinated animals. Similar to what has been shown in humans, COVID-19 after breakthrough infections with VoCs in vaccinated individuals have also been reported to be milder compared to infection in naïve, unvaccinated individuals^6, 40^.

It is well established that nAbs correlate with prevention of SARS-CoV-2 infection and reduced risk of severe disease^41^. Therefore, mean levels of nAbs elicited by different vaccines are generally predictive of their efficacy against SARS-CoV-2 infection and its variants^34, 41^. However, vaccine-induced nAb titers for VoCs can be much lower, in some cases >10-fold, than responses to the ancestral virus matching the original vaccine formulation deployed during the COVID-19 pandemic^42, 43^.

On the other hand, other groups have shown that there is very limited loss of T cell cross reactivity to VoCs, even genetically distant Omicron lineages^44, 45^. In accordance with these findings, our research shows that even in the absence of detectable titers of nAbs after suboptimal vaccination, vaccination shows a strong correlation with the T cell activation in the spleen upon infection with different VoCs, except for Beta. This suggests that the single mRNA vaccination already primed cross-reactive S-specific T cell responses. T cell boosting during Beta infection may have been low for several reasons. It is currently not known if S-specific immunodominant T cell epitopes in hamsters are affected by the mutations in the Beta S protein. Results in other animal models, including macaques for the Beta variant^46^ or humanized mouse models (K18-hACE2) for Gamma and Omicron variants^47^, also show that in the absence of nAbs or the presence of reduced nAbs titers, T cells control viral replication, disease, and lethality. The dissociation between nAbs and T cell responses could also play an important role in the control of infections with different vaccination strategies. While boosters could offer a much more robust nAb repertoire, as antibodies wane over time, T cells could play a critical role in protection against SARS-CoV-2, especially in regions where access to and deployment of repeated mRNA boosters is limited.

Our results show that hamsters vaccinated even with a suboptimal dose of BNT162b2 mount cross-reactive immune responses against antigenically distant variants. Early in the pandemic it was established that SARS-CoV-2 infection could protect against reinfection^29^. We observe in hamsters that virus challenge by itself generates a nAb response against the variant used for challenge, however, those nAb titers are higher in vaccine-primed animals as a result of hybrid immunity, similar to observations in humans^48, 49^. Infection in naïve unvaccinated hamsters resulted in cross-neutralizating antibodies upon infection with ancestral virus. However, the extent of infection-induced cross-neutralization was limited only to the antigenically closer variants such as Alpha and, to a lesser degree Delta, but was abrogated for the more distant Beta and Mu VoCs. A single BNT162b2 vaccination resulted in priming of cross-neutralizing antibody responses which, although undetectable at the moment of challenge, were boosted by infection by both vaccine-matching ancestral virus and more drifted VoCs. The improved cross-reactivity in post-challenge sera from vaccinated hamsters compared to that of unvaccinated animals is further illustrated by the smaller antigenic distances observed in antigenic cartography. Interestingly, challenge with 10^4^ PFU proved to be more efficient in the generation of nAbs against WA1/2020 than a single dose of BNT162b2. Broad neutralization against different VoCs after BNT162b2 vaccination has been shown in humans, although three homologous doses have been needed to induce potent nAbs against variants such as Beta or Omicron^50^.

Pathology upon SARS-CoV-2 infection in Syrian golden hamsters resemble those found in humans with mild SARS-CoV-2 infections^51^. Similar lung pathology has been reported in hamsters infected with different VoCs, with the exception of Omicron lineages, which show both lower viral loads in the lung as well as reduced pathology^52^. Our results show mild pathology both in vaccinated and unvaccinated hamsters except for the vaccinated group challenged with WA1/2020. Some pathology scores, particularly for the vaccinated group challenged with Alpha, might be driven by the reparative responses such as type II pneumocyte hyperplasia. Type II hyperplasia has been correlated with lung epithelial repair in Syrian golden hamsters after SARS-CoV-2 challenge^53^.

Gene expression patterns in the lung of infected animals correlate with prior findings of enhanced B and T cell activation upon infection in vaccinated animals. The effect of vaccination is greater in those groups that present improved protection (WA1/2020 and Alpha challenged), in line with the better antigenic match between challenge virus and vaccine. In groups that show lower protection expression of defense genes such as *Ddx60*, *Irf7*, *Rsad2*, *Ddx58*, *Stat2*, *Oas2*, *Ifitm2*, show an important role in the innate immune response and inflammatory responses, via RIGI and MDA5 mediated type I interferon responses^54–59^. Pro-inflammatory cytokines, such as *Cxcl10* also show higher activation in those groups less protected by the vaccine. Other genes involved in viral innate immune response show a similar activation pattern, such *Isg15*, which encodes a ubiquitin-like protein ISG15 that plays a key role in the innate immune response to viral infection either via its conjugation to a target protein (ISGylation) or via its action as a free or unconjugated protein^60^. Upregulation of genetic defense programs, pro-inflammatory and interferon-driven host responses in vaccinated animals when challenged with more antigenically distant VoCs is in line with lower control of virus replication and a stronger subsequent induction of the host immune response in these animals. Vaccination in animals challenged with the genetically-matched WA1/2020 shows to induce T-cell activation and B-cell proliferation and maturation through the upregulation of genes such as *Zfp683*, *Klrg1*, *Zc3h12d, Icosl*, *Lime1*, *Cd37*; as well as the recruitment of macrophages (*Zc3h12d*) and monocytes (*Nup85*).

Despite being a good model for SARS-CoV-2 infection studies, the Syrian golden hamster model comes with limitations when performing immune profiling of the host response to vaccination and infection due to the limited availability of antibodies that recognize immune markers. Therefore, we extrapolated immune cell abundances based on the host transcriptome using the Cibersortx algorithm. A drawback of this method, and all those relying on bulk RNA-seq data, is that cell abundances are relative and absolute numbers cannot be retrieved. This means that changes in abundance for one immune cell type is relative to that of other cell types and therefore need to be interpreted with caution. Moreover, the predicted immune cell abundances were obtained from RNA extracted from whole lung without perfusion. Therefore, we cannot make a difference between lung-resident immune cells like alveolar macrophages, circulating immune cells in blood and infiltrating immune cells like monocytic macrophages (M1), neutrophils, etc. Albeit these limitations, some useful observations were made. Alveolar macrophages are tissue resident myeloid immune cells and are a first immune barrier against pathogens in the alveolar space. Alveolar macrophages show reduced abundances in unvaccinated animals upon challenge. We and others have observed similar events in mice after experimental infection with influenza virus^61^. In the influenza mouse model, alveolar macrophages are transiently depleted in unvaccinated animals after infection and replenished after virus has been cleared from the lungs. This is typically associated with the infiltration of inflammatory monocytes, some of which become tissue-resident. The higher abundance of M1 macrophages in unvaccinated animals may reflect this phenomenon. Vaccination also results in lower levels of neutrophils compared to unvaccinated animals, further illustrating the protective effect of vaccination against ancestral virus as well as VoCs^62, 63^. Overall reduction in alveolar macrophages in the lung, as well as neutrophilia, are clearly shown in unvaccinated groups challenged with WA1/2020 and Alpha when compared to their respective vaccinated group, have been correlated with severe presentations of COVID-19 in humans^64^.

Interestingly, eosinophils are suggested to be enriched in vaccinated animals upon challenge with the antigenically matching WA1/2020 or Alpha variant. Pulmonary eosinophilia typically is correlated with a negative outcome of infection in vaccinated individuals, fueled by the vaccine associated enhancement of respiratory disease (VAERD) initially described for respiratory syncytial virus^65, 66^. However, antiviral effects of eosinophils have recently been suggested for both mice and humans^67, 68^, and we have recently described vaccine-associated pulmonary infiltration of eosinophils in the influenza mouse model that rather correlates with protection in the absence of VAERD^61^. In hamsters is associated with the orchestrated action of hyperstimulated macrophages and Th2 cytokine-secreting lymphoid cells^69^. Finally, frequencies for plasma cells were higher in vaccinated animals while T cells were higher in infected animals compared to mock challenged animals, regardless of their vaccination status. The different T cell activation profile in the lung by Cibersortx extrapolation when compared to the spleen by ELISPOT may be linked to the higher abundance of T memory cells in the spleen reactivated during infection when compared to the lung at this time point in infection^70^. In all, results are in line with the observed B and T cell responses further suggesting that mRNA vaccination, even when suboptimal, can prime for these adaptive immune responses and are boosted to superior levels (hybrid immunity) early on after infection both with ancestral vaccine-matching virus or more antigenically distant VoCs. In conclusion, we present a preclinical animal model to study host immune responses during breakthrough infections with ancestral SARS-CoV-2 and VoCs in mRNA vaccinated Syrian golden hamsters. We show that suboptimal vaccination is still protective, in the absence of virus-neutralizing titers at the moment of infection and skews host immune responses to infection. Hybrid immunity is the result of suboptimal vaccination followed by infection and results in cross-reactive B and T cell responses. These results illustrate that vaccination can contribute to protection with mechanisms beyond virus neutralization and therefore other immune correlates of protection than nAbs should be considered when evaluating vaccine responses.

## MATERIALS AND METHODS

### Key resources table

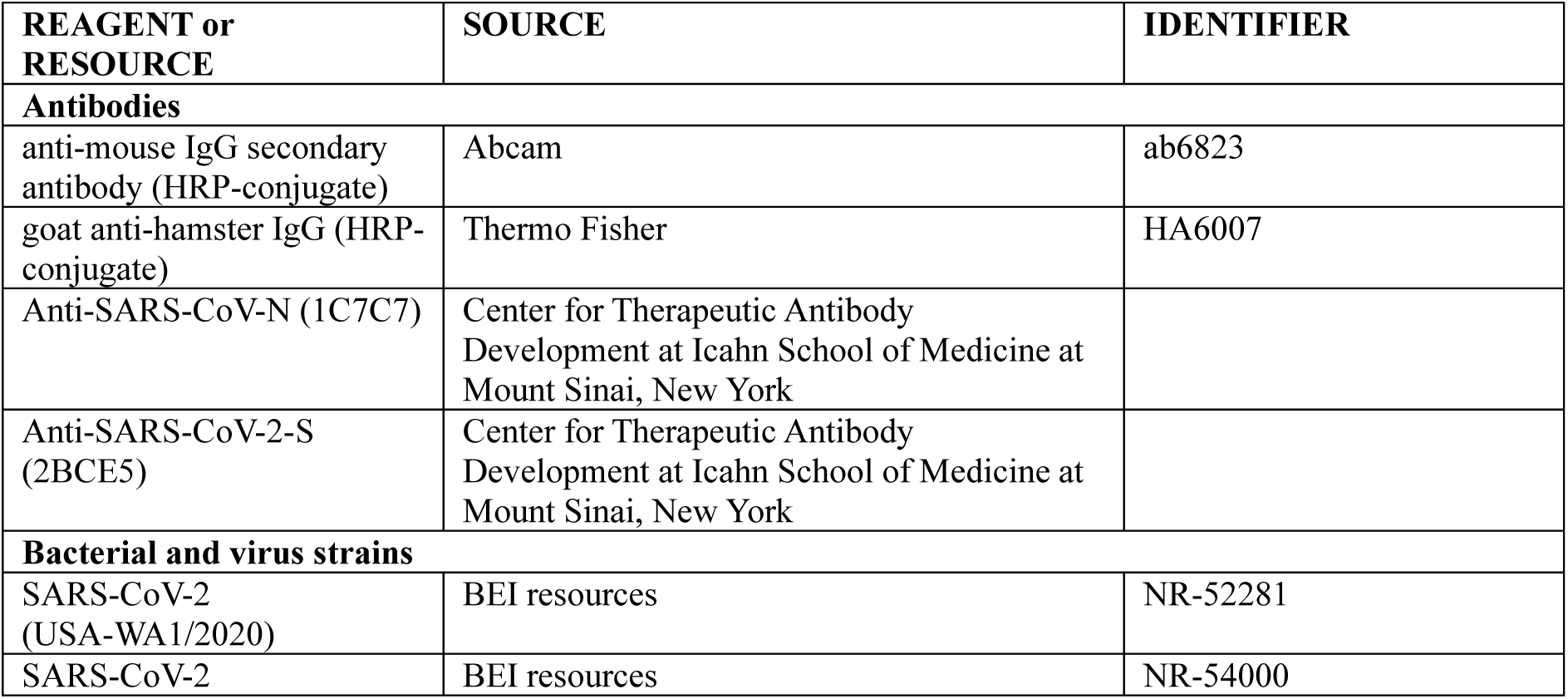
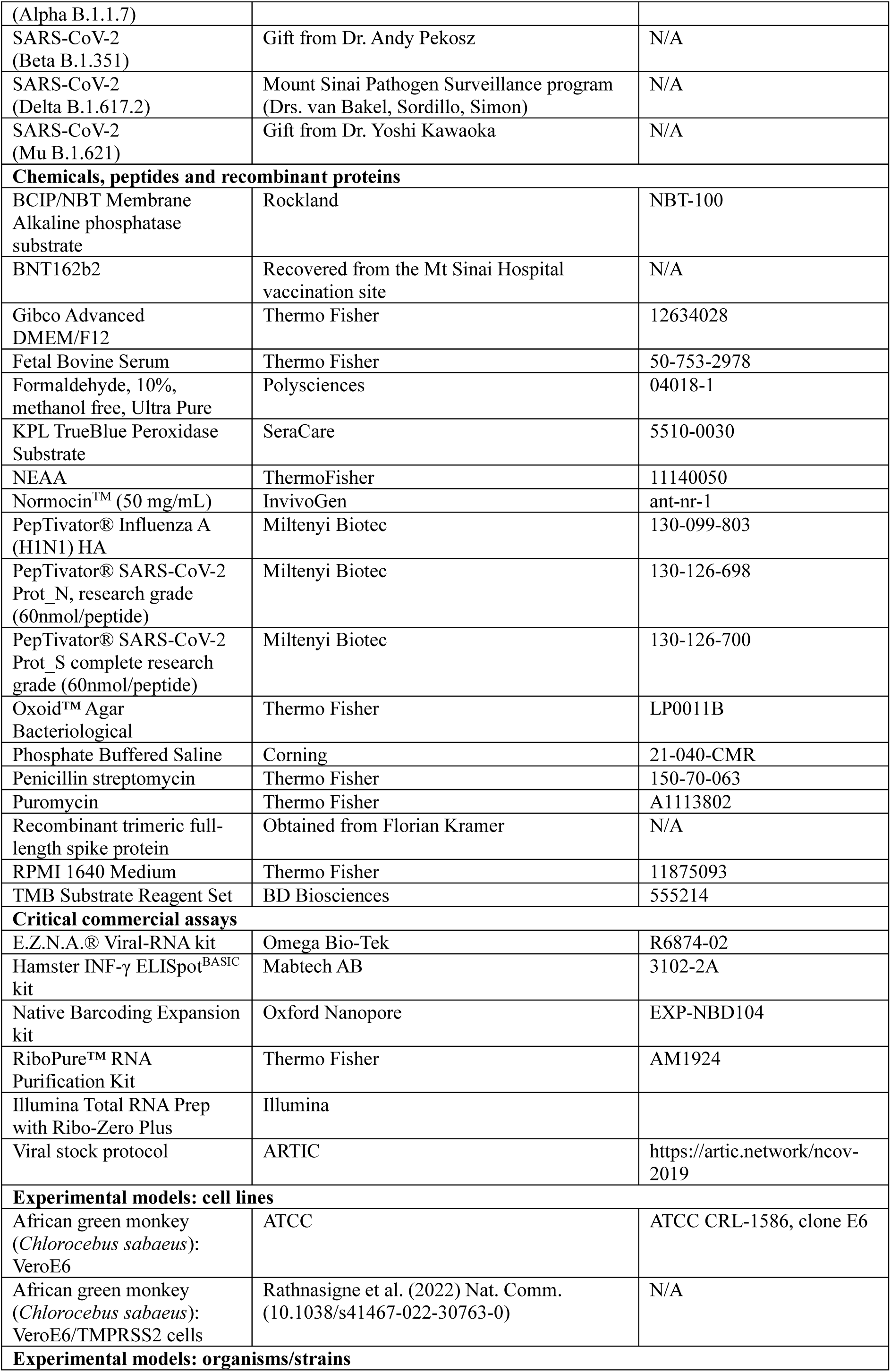
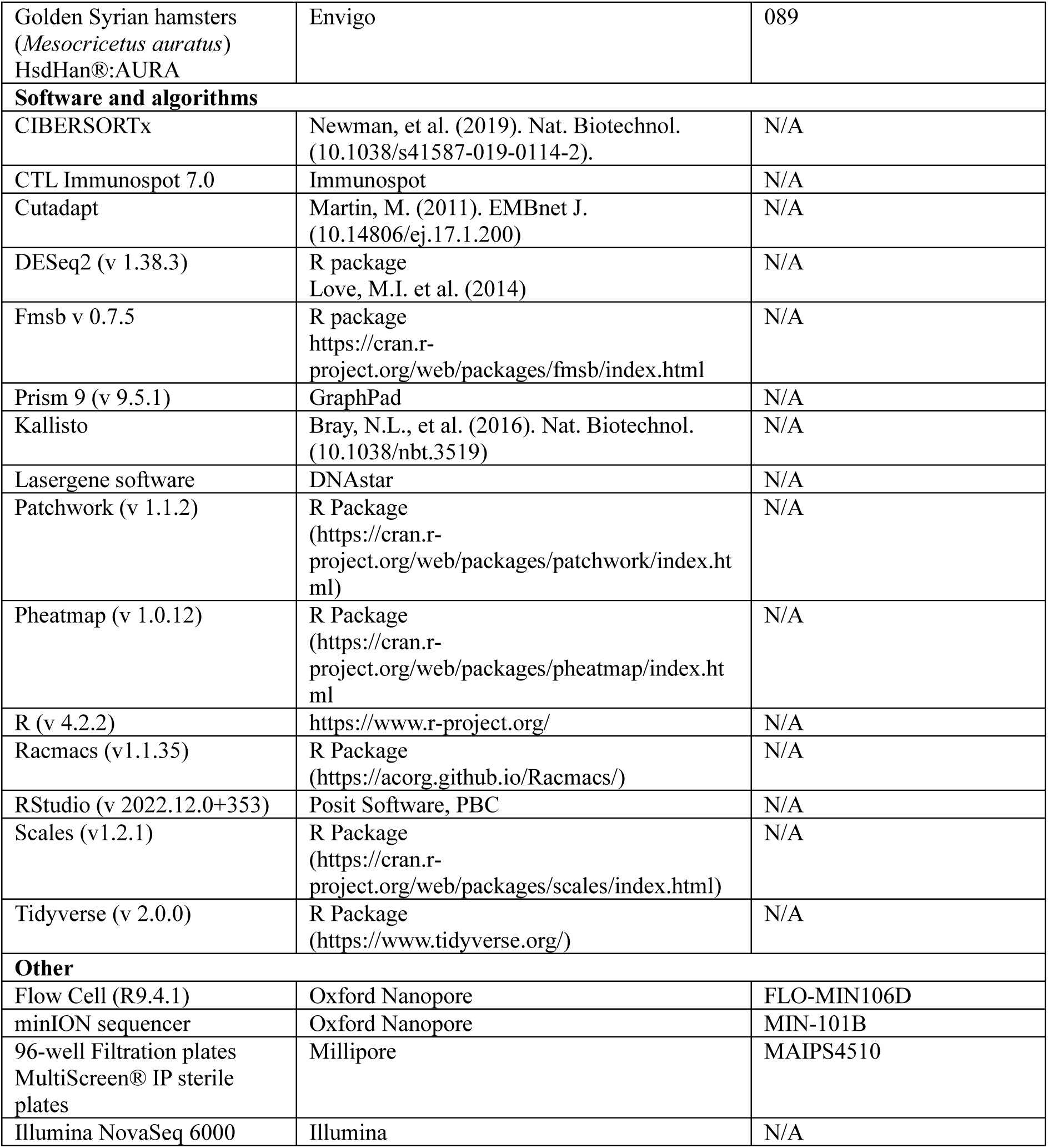

All experiments were approved and carried out in compliance with the Institutional Biosafety Committee (IBC) and Institutional Animal Care and Use Committee (IACUC) regulations of Icahn School of Medicine at Mt. Sinai.

### Cells

Vero-E6 cells (ATCC-CRL 1586, clone E6) were maintained in Dulbecco’s Modified Eagle’s Medium (DMEM; Gibco) containing 10% (v/v) of fetal bovine serum (FBS, Hyclone), 100 unit/mL of penicillin, 100 µg/mL of streptomycin (Gibco) and 1X of non essential amino acids (NEAA, Gibco). Vero-E6/TMPRSS2 cells, were transfected to stabily express the serine protease TMPRSS2 under a puromycin selection marker. They were maintained in DMEM containing 10% (v/v) FBS, 100 unit/mL of penicillin, 100 µg/mL of streptomycin, 1% NEAA, 3 µg/mL of puromycin and 100 µg/mL of normocin.

### Viruses

SARS-CoV-2 isolate USA-WA1/2020 was obtained from BEI resources (NR-52281) and was propagated on Vero-E6 cells as previously described (33073694). Alpha variant (B.1.1.7, hCoV-19/England/204820464/2020) was obtained from BEI Resources (NR-54000). Beta variant (B.1.351, hCoV-19/USA/MD-HP01542/2021 JHU) was a kind gift from Dr. Andy Pekosz. Delta variant (B.1.617.2; hCoV-19/USA/NYMSHSPSP-PV29995/2021) is a clinical isolate provided by the Mount Sinai Pathogen Surveillance Program, directed by Drs. van Bakel, Sordillo and Simon). Mu variant (B.1.621, hCoV-19/USA/WI-UW-4340/2021) was a kind gift from Dr. Yoshi Kawaoka. Alpha, Beta, Delta and Mu variants were propagated on VeroE6/TMPRSS2. All stocks were titrated on VeroE6/TMPRSS2 cells by plaque assay and 50% tissue culture infective dose (TCID50).

### Deep sequencing of the viral stocks

All viral stocks were sequence-confirmed using the ARTIC protocol (https://artic.network/ncov-2019, Primer set version 3). Briefly, viral RNA was purified using E.Z.N.A.^®^ Viral-RNA kit (Omega-Bio-Tek) according to the manufacturer’s instructions and was used to prepare cDNA. Overlapping amplicons of ∼400 bp covering the whole genome were barcoded using the Oxford Nanopore Technologies Native Barcoding Expansion kit (EXPNBD104). Libraries were prepared according to the manufacturer’s instructions and loaded on a minION sequencer equipped with a FLO-MIN106D flow cell. The consensus sequence was obtained using Lasergene software (DNAstar).

### Hamster immunization and challenge

Sixty 12 to 14-week-old female Golden Syrian hamsters (*Mesocricetus auratus*) were procured from the Envigo vendor and housed in specified pathogen-free conditions. On 0 days post-vaccination (0 DPV), thirty hamsters were intramuscularly immunized with a single 5 µg dose (100 µL) of Pfizer mRNA vaccine, BNT162b2 (vaccinated group), and the other thirty were injected with 100 µL of 1X Phosphate buffer saline (PBS) (unvaccinated group). BNT162b2 vaccine was recovered from the Mt Sinai Hospital vaccination site and stored frozen at -80°C after freeze-thaw. Vaccine immunogenicity was confirmed by dose titration experiments in mice using ELISA, virus microneutralization and T cell assays as read out.

On 21 DPV, blood was collected from all hamsters from vena-cava under mild ketamin/xylasin anesthesia. For the SARS-CoV-2 virus challenge, all hamsters were transferred to an Animal Biosafety level 3 (ABSL3) facility two days prior to the viral challenge. On 28 DPV, 25 of the anesthetized hamsters from the vaccinated or unvaccinated groups were intranasally challenged with 1×10^4^ plaque forming units (PFU, per hamster in 100 µL of volume) of either USA/WA1/2020 (WA1/2020, ancestral), Alpha, Beta, Delta, or Mu variants of SARS-CoV-2 (N=5/group), rest of 35 hamsters from both vaccinated or unvaccinated groups were given 100 µL of 1X PBS as mock infection under anesthesia. Animals were maintained up to 5 days post-infection (DPI) and were monitored daily for body weight changes. On 5 DPI (32 DPV), all hamsters were euthanized, blood was collected via terminal cardiac puncture, and the spleen and lungs were collected for further analysis.

### Enzyme-linked Immunosorbent Assay (ELISA)

SARS-CoV-2 spike-specific ELISA assays were performed as previously described^71^. In brief, Nunc MaxiSorp^TM^ flat-bottom 96-well plates (Invitrogen) were coated with 2µg/mL of recombinant soluble trimeric full-length spike protein (50 µL per well, produced in HEK293T cells using pCAGGS plasmid vector containing Wuhan-Hu-1 Spike glycoprotein gene produced under HHSN272201400008C and obtained through BEI Resources, NR-52394) in bicarbonate buffer overnight at 4°C. Plates were then washed three times with 1x PBST (1x PBS + 0.1% v/v Tween-20). Then, plates were blocked with 100 µL per well of blocking solution (5% non-fat dry milk in PBST) for 1 hr at room temperature (RT). The blocking buffer was decanted, and hamster sera were threefold serially diluted in the blocking solution starting at 1:100 dilution and incubated for 1.5 h RT. The plates were washed three times with in PBST and 50 µL of HRP-conjugated goat anti-hamster IgG (H+L) cross-adsorbed secondary antibody (Invitrogen, HA6007) was added at 1:5000 dilution. The plates were incubated for 1 hr at RT and washed 3 times with PBST. Finally, 100 μL tetramethyl benzidine (TMB; BD optiea) substrate was added and incubated at RT until blue color was developed. The reaction was stopped with 50μl 1M H2SO4 and absorbance was recorded at 450nm and 650nm. An average of OD450 values for blank wells plus three standard deviations was used to set a cutoff value for each plate.

### 50% tissue culture infective dose (TCID50) and *in vitro* microneutralization assays

Estimation of neutralization capacity of sera from vaccinated and unvaccinated hamsters, both pre and post-challenge, was performed by *in vitro* microneutralization assays, based on previously described methodology^72^. First, for TCID50 calculation, the viral stocks were serially diluted 10-fold starting with 1:10 dilution in a total volume of 100 µL and incubated on Vero-E6/TMPRSS2 cells for 48 hours followed by fixation in 4% methanol free formaldehyde. For immunostaining, the cells were washed with 1x PBST and incubated in 100 µL permeabilization buffer (0.1% Triton X-100 in 1x PBS) for 15 min at RT. The cells were washed again with 1x PBST and blocked in 5% non-fat milk in 1x PBS-T for 1 hr at RT. After blocking, the cells were incubated with 1:1000 of anti-SARS-CoV-2-N and anti-SARS-CoV-2-Spike monoclonal antibodies, for 1.5 hr at RT. Subsequently, the cells were washed in 1x PBST and incubated with 1:5000 diluted HRP conjugated anti-mouse IgG secondary antibody (ab6823) for 1 hr at RT followed by another PBST wash. Finally, 100μl TMB substrate was added and incubated at RT until blue color was developed. The reaction was stopped with 50μl 1M H2SO4 and absorbance was recorded at 450 nm and 650 nm.

For *in-vitro* micro-neutralization assays, the serum samples were inactivated at 56°C for 30 min. The sera were serially diluted 3-fold starting from 1:20 dilution in infection medium (DMEM, 2% FBS and 1% NEAA). Sera dilutions were incubated with 350TCID50/well of USA-WA1/2020, Alpha, Beta, Delta and Mu variants, for 1 hour in an incubator at 37°C and 5% CO2; followed by incubation on pre-seeded Vero E6/TMPRSS2 at 37°C, 5% CO2 for 48 hours. The plates were fixed in 4% formaldehyde followed by immunostaining, similar to TCID50 assays. Percentage of neutralization was calculated in reference to the mean of negative and positive controls for each plate and half of the maximum inhibitory dose (ID50) was calculated for each serum sample against individual viruses using GraphPad prism.

### Virus Challenge

1×10^4^ plaque forming units (PFU) per animal of each of USA-WA1/2020 (ancestral), Alpha, Beta, Delta or Mu variants was used for intranasal infection, in a final volume of 100 µL per animal, performed under deep ketamine/xylazine sedation. Five vaccinated and five unvaccinated animals were mock-challenged and referred to as mock groups for reference in the study. Therefore, 12 animal groups (5 animals/ group) were established: PBS-mock, BNT162b2-mock, PBS-WA1/2020, BNT162b2-WA1/2020, PBS-Alpha, BNT162b2-Alpha, PBS-Beta, BNT162b2-Beta, PBS-Delta, BNT162b2-Delta, PBS-Mu, BNT162b2-Mu. Body weights were recorded every day to assess the morbidity post infection until organ harvest. Terminal blood collection was done by the cardiac puncture, along with lungs and spleen harvest *post-mortem* at 5 DPI.

### Virus titration by plaque assays

Plaque assays were performed to determine viral titers in lungs from hamsters challenged with USA-WA1/2020, Alpha, Beta, Delta or Mu variants. Briefly, lungs were harvested from the animals and the cranial and middle lobes from the right lung were collected and homogenized in sterile 1x PBS for viral titration. Centrifugation was performed at 7,000 g for 5 minutes to remove tissue debris. Then, the homogenates were 10-fold serially diluted starting from a 1:10 dilution in 1x PBS. Pre-seeded Vero-E6/TMPRSS2 cells were infected with diluted lung homogenates for 1 hour at RT with occasional shaking, followed by an overlay of 2% oxoid agar mixed with 2x MEM supplemented with 0.3% FBS. The plates were incubated for 72 hours at 37°C and 5% CO2 followed by fixation in 1mL of 10% methanol-free formaldehyde. The plaques were immuno-stained with anti-mouse SARS-CoV-2-N antibody diluted 1:1000 in 1x PBST for 1.5 hr at RT with gentle shaking and subsequently with 1:5000 diluted HRP-conjugated anti-mouse secondary IgG antibody for 1 hr at RT. Finally, the plaques were developed with KPL TrueBlue Peroxidase Substrate (Seracare) and viral titers were calculated and represented as plaque forming units (PFU)/mL.

### Antigen cartography

Antigenic maps were constructed as previously described^73^. In brief, antigenic cartography allows quantification and visualization of neutralization data by computing distances between antiserum and antigen points. Distances correspond to the difference between the log2 of the maximum titer observed for an antiserum against any antigen and the log2 of the titer for the antiserum against a specific antigen. Modified multidimensional scaling methods are then used to arrange the antigen and antiserum points in an antigenic map. In the resulting map, the distance between points represents antigenic distance as measured by the neutralization assay, in which the distances between antigens and antisera are inversely related to the log2 titer^74^. Maps were computed with the R package “Racmacs” (https://acorg.github.io/Racmacs/, version 1.1.35.), using 1000 optimizations, with the minimum column basis parameter set to “none”. Sera yielding microneutralization results under the limit of detection were set to 10 for all calculations.

### IFN**-**γ^+^ ELISPOTs

Spleens were harvested from all hamsters and collected in RPMI-1640 media supplemented with 10% FBS and 1x Penicillin/Streptomycin. Single cell splenocyte suspensions were prepared by forcing the spleens through a 70 μm cell strainer. IFN-γ^+^ ELISPOTs assays were performed using 10^5^ cells/well with the Hamster IFN-γ ELISPOT^BASIC^ kit (Mabtech AB) following manufacturer’s instructions. In brief, 96-well Millipore polyvinylidene difluoride (PVDF) plates were treated with 50 μL of 70% ethanol for 2 minutes, then thoroughly washed 5 times with sterile water (200 μL/well). Plates were coated with 100 μL of 15 μg/mL coating antibody (clone H21) and left overnight at 4°C. Plates were washed five times and then 200 μL/well of medium was added to the plate for 30 min at room temperature. After removing the medium, the splenocyte suspension (10^5^ cells/well) was added to the plate and splenocytes were stimulated overnight at 37°C and 5% CO2 with Spike, Nucleoprotein or Hemagglutinin overlapping peptide pools (Miltenyi Biotec: PepTivator® SARS-CoV-2 Prot_S complete, PepTivator® SARS-CoV-2 Prot_N, PepTivator® Influenza A (H1N1) HA, respectively). Medium was then removed, wells were washed thoroughly and cells were fixed with 4% methanol-free formaldehyde before transferring out from BSL-3 facility, and plates were thoroughly washed in PBS (200 μL/well). Then, 100 μL/well of 1 μg/mL of detection antibody (H29-biotin) diluted in PBS supplemented with 0.5% FBS were added to the plate, which was incubated for 2h at RT. After washing with PBS, Streptavidin-ALP diluted in PBS-0.5% FBS (1:1000) and incubated for 1h at RT. Once more, plates were washed with PBS and 100 μL/well of BCIP/NBT substrate solution were added and developed until color emerged. Finally, the plates were thoroughly washed under tap water 5 time and allowed to air dry overnight in dark. The number of spots in each well were represented as number of IFNγ producing cells per million splenocytes or fold induction of S or N stimulated cells over unspecific stimulation of cells with HA.

### Histopathology

On the day of necropsy, left lungs were inflated with 4% formaldehyde in PBS and fixed. Lungs were sectioned longitudinally on the midline to exposed central airways and vessels. Both portions were routinely processed, embedded in paraffin blocks and stained with hematoxylin-eosin (H&E). The slides were assessed blinded to treatment and evaluated randomly. A pathological scoring system was used to assess ten parameters: amount of lung affected, perivascular inflammation, vasculitis/ vessel injury, bronchial/bronchiolar necrosis, bronchial/bronchiolar inflammation, alveolar inflammation, alveolar necrosis, alveolar edema and type II pneumocyte hyperplasia. Histological parameters were scored in a scale of 0 to 5. In terms of area affected a score of 0 indicated none, 1=5-10%, 2=10-25%, 3=25-50%, 4=50-75% and 5>75%. For histological parameters: 0=none, 1=minimal, 2=mild, 3=moderate, 4=marked, 5=severe. Overall lesion scores were also calculated and scaled to 0-5 for representation. Radar charts were generated with histopathological scores in R, using “fmsb” (v.0.7.5) and “scales” (v.1.2.1) packages.

### RNA-seq

Right lung caudal lobe was harvested from the animals and homogenized in trizol. RNA purification was performed with the kit RiboPure™ RNA Purification Kit following manufacturer’s instructions. Sequencing libraries were generated with the Illumina Total RNA Prep with Ribo-Zero Plus kit. These libraries were sequenced on an Illumina NovaSeq 6000 using an S4 flow cell, generating paired-end 100 bp reads. Adapter sequences were trimmed from reads with Cutadapt^75^, then pseudoaligned by Kallisto^76^ to the Golden Hamster transcriptome. The transcriptome index was built from the MesAur1.0 genome assembly and gene annotation from Ensembl release 100. Differentially expressed genes were identified using DESeq2^77^. Cell types were deconvoluted from bulk RNA-seq profiles with CIBERSORTx^78^. A signature matrix was built from a single-cell Golden Hamster lung dataset^31^, which was randomly down sampled to 100 cells of each type. Cell fractions were imputed using default parameters and a normalized count matrix as input. Analysis were performed and graphs were performed in R, using the packages “DESeq2” (v 1.38.3), “pheatmap” (v 1.0.12) and “patchwork” (v 1.1.2).

### Other statistical analysis

Comparisons between vaccinated and unvaccinated groups were performed by Mann Whitney U test. They were performed with GraphPad Prism 9 (GraphPad Software). * represents p-value ≤ 0.05; ** represents p-value ≤ 0.01; *** represents p-value ≤ 0.001 and **** represents p-value ≤ 0.0001.

## ACKNOWLEDGEMENTS

We thank Daniel Flores, Marlene Espinoza, Jane Deng, and Ryan Camping for excellent administrative support, Richard Cadagan for technical support and Randy Albrecht for management and organization of the BSL3 facility. We want to thank all colleagues from the NIH SAVE consortium for their input and feedback. This work was partially supported by CRIPT (Center for Research on Influenza Pathogenesis and Transmission), a NIAID Center of Excellence for Influenza Research and Response (CEIRR, contract # 75N93021C00014) and by NIAID grant U19AI135972 to AG-S. SARS-CoV-2 work in the M.S. laboratory is supported by NIH/NIAID R01AI160706 and NIH/NIDDK R01DK130425. K.H. acknowledges the <u>M.Sc</u>. program Integrated Immunology Immunology (supported by the Elite Network of Bavaria) at the Friedrich Alexander University Erlangen-Nuremberg. This publication includes data generated at the UC San Diego IGM Genomics Center utilizing an Illumina NovaSeq 6000 that was purchased with funding from a National Institutes of Health SIG grant (#S10 OD026929).

