## Supplementary Table for "Breakthrough infections by SARS-CoV-2 variants boost cross-reactive hybrid immune responses in mRNA-vaccinated Golden Syrian Hamsters"

### Supplementary Table 1. Genes upregulated by vaccination but not by infection in vaccine-matching WA1/2020 infected animals.

| Ensembl Code | Gene | Base mean[a] | $\log_2$ (Fold Change)[b] | lfcSE[c] | Stat[d] | p-value | p-adj[e] |
|---|---|---|---|---|---|---|---|
| **ENSMAUG00000007886** | *Vsig1* | 23.82840 | 2.82632 | 0.86353 | 3.27297 | 0.00106 | 0.02395 |
| **ENSMAUG00000020451** | *Zfp683* | 12.77957 | 2.66024 | 0.47628 | 5.58548 | 0.00000 | 0.00000 |
| **ENSMAUG00000005020** | *Hrob* | 8.07341 | 2.60012 | 0.71668 | 3.62798 | 0.00029 | 0.00896 |
| **ENSMAUG00000014751** | *Itln1* | 72.86937 | 2.23639 | 0.63190 | 3.53918 | 0.00040 | 0.01157 |
| **ENSMAUG00000001649** | *Cyp26b1* | 26.26286 | 2.05161 | 0.62028 | 3.30753 | 0.00094 | 0.02209 |
| **ENSMAUG00000001547** | *Spns3* | 12.61973 | 1.97350 | 0.45548 | 4.33284 | 0.00001 | 0.00079 |
| **ENSMAUG00000000757** | *Dbp* | 28.13340 | 1.72906 | 0.49749 | 3.47559 | 0.00051 | 0.01379 |
| **ENSMAUG00000015676** | *Kcng1* | 8.66631 | 1.71226 | 0.48195 | 3.55276 | 0.00038 | 0.01112 |
| **ENSMAUG00000015196** | *Sh2d1a* | 10.36465 | 1.67394 | 0.54749 | 3.05750 | 0.00223 | 0.04050 |
| **ENSMAUG00000015783** | *Zc3h12d* | 19.91885 | 1.57167 | 0.47552 | 3.30513 | 0.00095 | 0.02224 |
| **ENSMAUG00000017156** | *Marchf10* | 7.06600 | 1.55809 | 0.50802 | 3.06698 | 0.00216 | 0.03966 |
| **ENSMAUG00000018293** | *Arhgef39* | 10.35627 | 1.53928 | 0.51236 | 3.00431 | 0.00266 | 0.04591 |
| **ENSMAUG00000015031** | *Klrg1* | 9.36680 | 1.39950 | 0.40779 | 3.43188 | 0.00060 | 0.01552 |
| **ENSMAUG00000019266** | *Haao* | 14.74745 | 1.39186 | 0.39352 | 3.53694 | 0.00040 | 0.01160 |
| **ENSMAUG00000021348** | *Col11a2* | 17.03576 | 1.37335 | 0.36844 | 3.72742 | 0.00019 | 0.00651 |
| **ENSMAUG00000019140** | *Icosl* | 18.74959 | 1.34756 | 0.38403 | 3.50895 | 0.00045 | 0.01250 |
| **ENSMAUG00000018778** | *Lime1* | 11.40338 | 1.28389 | 0.42094 | 3.05003 | 0.00229 | 0.04121 |
| **ENSMAUG00000017163** | *Fcho1* | 23.10064 | 1.08662 | 0.29468 | 3.68740 | 0.00023 | 0.00745 |
| **ENSMAUG00000000752** | *Ddx11* | 19.85041 | 1.03004 | 0.33604 | 3.06521 | 0.00218 | 0.03984 |
| **ENSMAUG00000022124** | *Wdr76* | 372.80547 | 1.00393 | 0.29214 | 3.43648 | 0.00059 | 0.01532 |
| **ENSMAUG00000014187** | *Sult1c2* | 21.36362 | 0.99204 | 0.29838 | 3.32474 | 0.00089 | 0.02107 |
| **ENSMAUG00000020400** | *Ctc1* | 25.34021 | 0.92440 | 0.27417 | 3.37159 | 0.00075 | 0.01846 |
| **ENSMAUG00000015203** | *Nt5c* | 55.46074 | 0.89023 | 0.24775 | 3.59328 | 0.00033 | 0.00988 |
| **ENSMAUG00000014228** | *Lrrc18* | 22.63506 | 0.88098 | 0.29365 | 3.00007 | 0.00270 | 0.04612 |
| **ENSMAUG00000007672** | *Ltb* | 61.80016 | 0.87800 | 0.26600 | 3.30068 | 0.00096 | 0.02230 |
| **ENSMAUG00000018364** | *Ifrd2* | 92.22219 | 0.87744 | 0.22490 | 3.90147 | 0.00010 | 0.00367 |
| **ENSMAUG00000016464** | *Rfc3* | 42.17511 | 0.86357 | 0.25933 | 3.33001 | 0.00087 | 0.02085 |
| **ENSMAUG00000010583** | *Cdca7l* | 71.94039 | 0.86094 | 0.26738 | 3.21986 | 0.00128 | 0.02696 |

| | | | | | | | |
|---|---|---|---|---|---|---|---|
| **ENSMAUG00000016884** | *Themis* | 36.94183 | 0.79483 | 0.25709 | 3.09161 | 0.00199 | 0.03717 |
| **ENSMAUG00000003857** | *Slirp* | 39.71779 | 0.78927 | 0.25485 | 3.09698 | 0.00196 | 0.03678 |
| **ENSMAUG00000020536** | *Nup85* | 62.99557 | 0.75629 | 0.19462 | 3.88605 | 0.00010 | 0.00386 |
| **ENSMAUG00000019979** | *Lmnb2* | 135.68315 | 0.74401 | 0.20230 | 3.67773 | 0.00024 | 0.00766 |
| **ENSMAUG00000020606** | *Srgap3* | 53.32324 | 0.72252 | 0.20753 | 3.48148 | 0.00050 | 0.01359 |
| **ENSMAUG00000018732** | *Tssc4* | 50.93902 | 0.64719 | 0.20294 | 3.18913 | 0.00143 | 0.02885 |
| **ENSMAUG00000012212** | *Cd37* | 114.90059 | 0.63391 | 0.20603 | 3.07673 | 0.00209 | 0.03877 |
| **ENSMAUG00000003325** | *Paxip1* | 93.22434 | 0.61247 | 0.14099 | 4.34399 | 0.00001 | 0.00076 |
| **ENSMAUG00000020021** | *Pfas* | 171.59328 | 0.60483 | 0.18879 | 3.20379 | 0.00136 | 0.02777 |
| **ENSMAUG00000012577** | *Lig1* | 186.52698 | 0.59510 | 0.16959 | 3.50904 | 0.00045 | 0.01250 |
| **ENSMAUG00000000104** | *Gpr183* | 109.25363 | 0.53348 | 0.17415 | 3.06331 | 0.00219 | 0.03997 |
| **ENSMAUG00000019805** | *Tarbp2* | 92.34949 | 0.48403 | 0.15314 | 3.16062 | 0.00157 | 0.03111 |
| **ENSMAUG00000009284** | *Stxbp2* | 160.02297 | 0.42772 | 0.13681 | 3.12635 | 0.00177 | 0.03426 |
| **ENSMAUG00000007062** | *Eef2* | 4806.03639 | 0.40648 | 0.12531 | 3.24379 | 0.00118 | 0.02536 |
| **ENSMAUG00000014174** | *Gpi1* | 418.74015 | 0.40584 | 0.11303 | 3.59054 | 0.00033 | 0.00994 |
| **ENSMAUG00000018966** | *Khsrp* | 306.17420 | 0.36374 | 0.11214 | 3.24356 | 0.00118 | 0.02536 |
| **ENSMAUG00000009823** | *Esyt1* | 472.96805 | 0.35912 | 0.10721 | 3.34982 | 0.00081 | 0.01968 |
| **ENSMAUG00000022241** | *Tp53* | 183.25934 | 0.34594 | 0.10841 | 3.19113 | 0.00142 | 0.02875 |

[a]Base: Mean average of the normalized count values, dividing by size factors, taken over all samples.
[b]$log_2$ (Fold Change): Effect size estimate.
[c]lfcSE: standard error estimate $log_2$ (Fold Change).
[d]stat: value of the test statistic for the transcript.
[e]p-adj: Adjusted p-value for multiple testing for the transcript.
