## Supplementary Figures for "Breakthrough infections by SARS-CoV-2 variants boost cross-reactive hybrid immune responses in mRNA-vaccinated Golden Syrian Hamsters"

**Supplementary Figure 1.** INF-y producing cells per million splenocytes after stimulation with (from left to right): N-peptide, S-peptide, HA-peptide and un-stimulated.

**Supplementary Figure 2. Gene ontology.** Top 25 most significantly upregulated biological processes based on significance in WA1/2020 challenged groups. (A) Upregulated biological processes in vaccinated animals compared to unvaccinated (B) Upregulated biological processes in unvaccinated animals compared to vaccinated.

**Supplementary Figure 3. Abundance of additional cell subsets in the lungs of infected hamsters extrapolated from the host transcriptome.** Scale represents estimated fraction of each cell type for each condition.

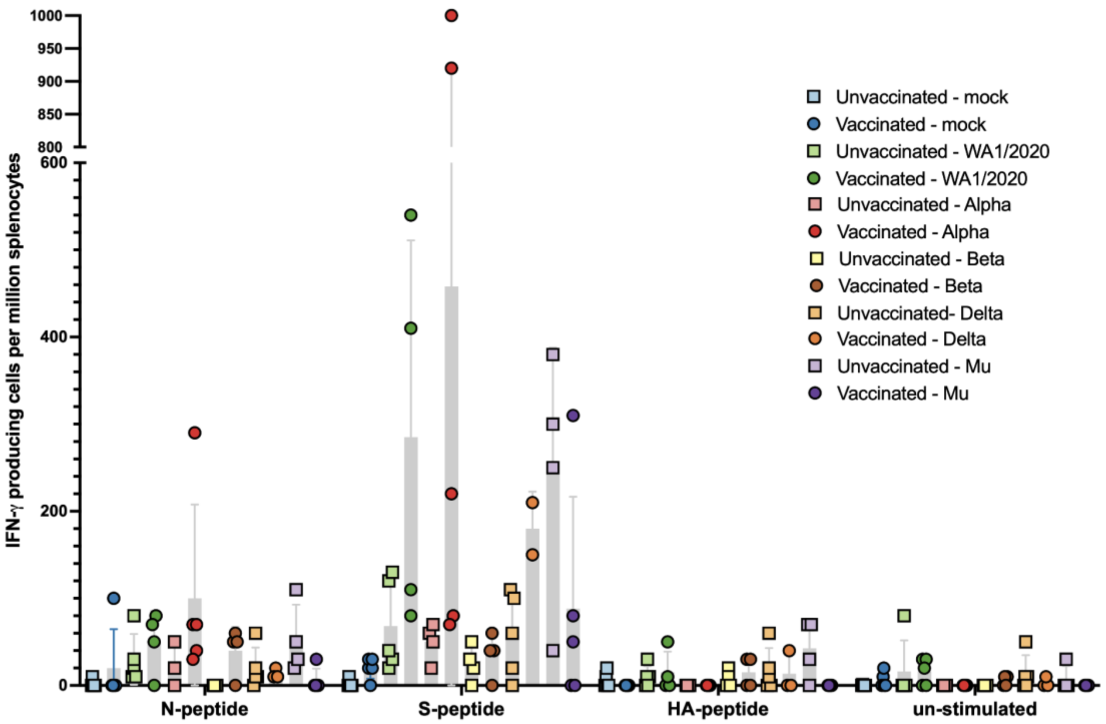


**Supplementary Figure 1.** INF-y producing cells per million splenocytes after stimulation with (from left to right): N-peptide, S-peptide, HA-peptide and un-stimulated.

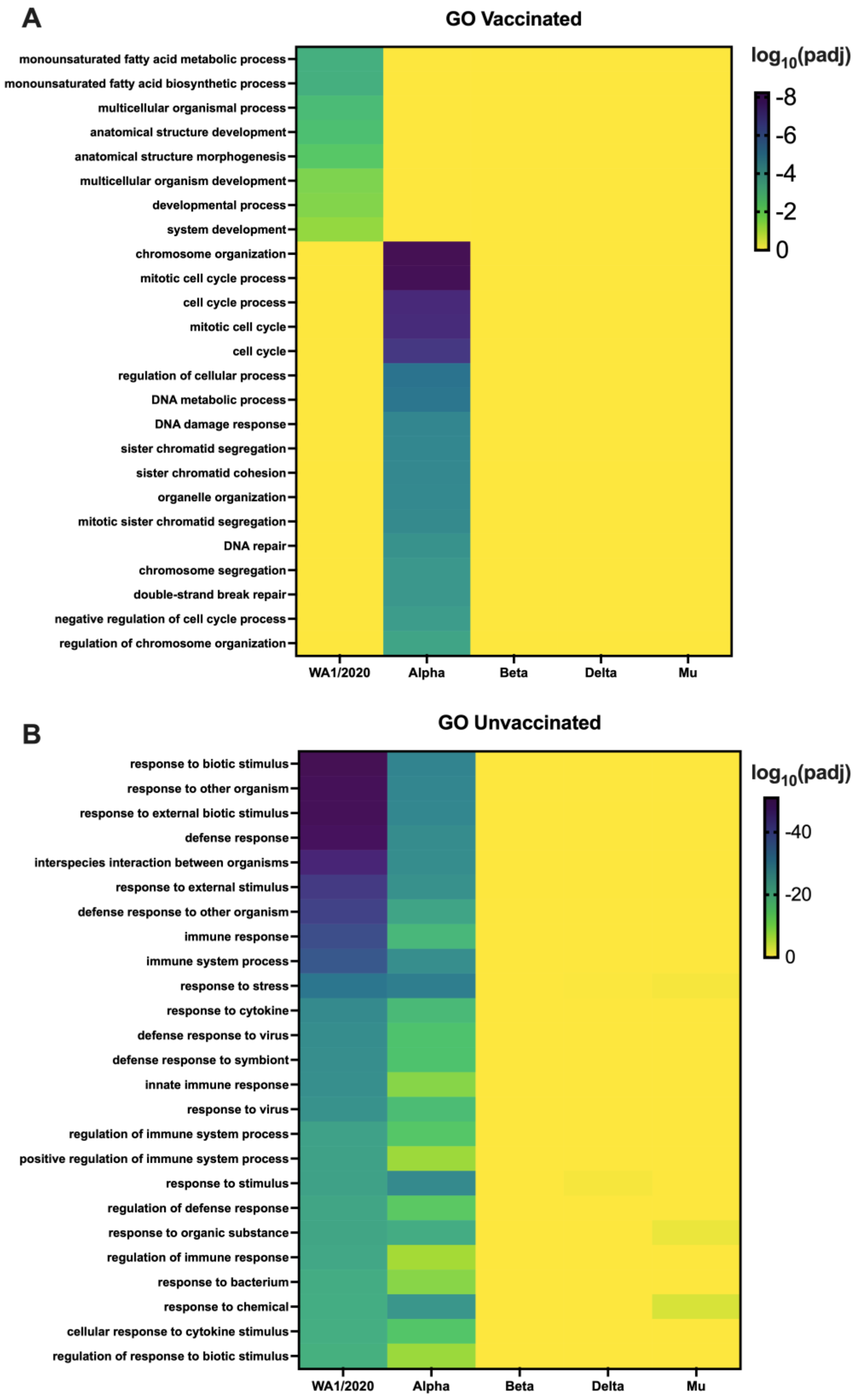


**Supplementary Figure 2. Gene ontology.** Top 25 most significantly upregulated biological processes based on significance in WA1/2020 challenged groups. (A) Upregulated biological processes in vaccinated animals compared to unvaccinated (B) Upregulated biological processes in unvaccinated animals compared to vaccinated.

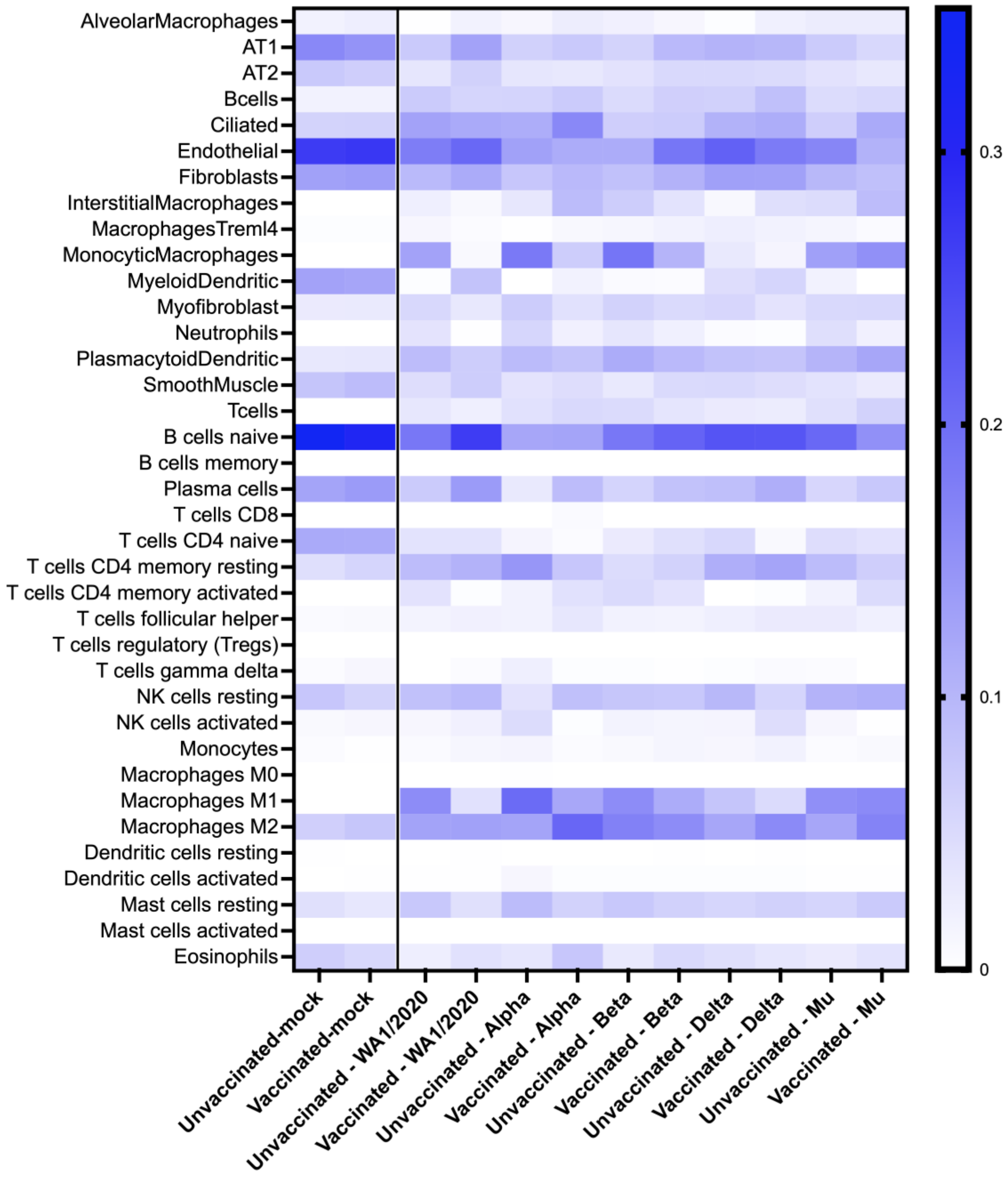


**Supplementary Figure 3. Abundance of additional cell subsets in the lungs of infected hamsters extrapolated from the host transcriptome.** Scale represents estimated fraction of each cell type for each condition.
